# Small Particles, Big Problems: Polystyrene nanoparticles induce DNA damage, oxidative stress, migration, and mitogenic pathways predominantly in non-malignant lung cells

**DOI:** 10.1101/2025.03.24.644975

**Authors:** Büsra Ernhofer, Andreas Spittler, Franziska Ferk, Miroslav Mišík, Martha Magdalena Zylka, Lisa Glatt, Kristiina Boettiger, Anna Solta, Dominik Kirchhofer, Gerald Timelthaler, Zsolt Megyesfalvi, Verena Kopatz, Heinrich Kovar, Siegfried Knasmueller, Clemens Aigner, Lukas Kenner, Balazs Döme, Karin Schelch

## Abstract

Polystyrene micro-and nanoplastics (PS-MNPs) are emerging environmental pollutants with potential implications for human health. In this study, we used two different sizes of PS-MNPs (0.25 µm and 1 µm) on non-small cell lung cancer (A549, H460), small cell lung cancer (DMS53, H372), and normal lung epithelial (BEAS-2B) cells, as well as on human-derived lung organoids, to investigate the cytotoxic effects of PS particles. At lower concentrations (< 30 µg/cm^2^, equivalent to 50 µg/ml), neither PS-MPs nor PS-NPs did not interfere with cell viability or proliferation. Intracellular kinetic assays revealed that non-malignant (BEAS-2B) lung cells showed the strongest turnover of PS-NPs compared to malignant cells. Since PS-NPs exhibited more pronounced cellular effects, we focused further analyses on their impact. Furthermore, we observed significantly increased migration, prolonged S-phase arrest along with induced DNA damage, and oxidative stress in non-malignant (BEAS-2B) lung cells. Thus, our data suggest that BEAS-2B cells exhibit the highest sensitivity to PS-NPs. We also demonstrate that after PS-NP treatment, these cells displayed decreased base excision repair capacity and increased activation of survival pathways, including AKT and ERK phosphorylation. PS-NP internalization and increase of signal pathways were validated in a more physiological lung organoid setting. Altogether, our findings suggest that PS-NPs do not significantly affect the malignant behavior of cancer cells. However, they could promote tumor-like features in normal lung cells by inducing survival pathways, migration, and alterations in stress response mechanisms.

**Environmental Implications:** This study investigates the effects of polystyrene micro-and nanoplastics (PS-MNPs) at environmentally relevant concentrations. The tested concentrations of PS-MNPs (0, 15, 30, and 60 µg/cm^2^, equivalent to 0, 25, 50, and 100 µg/ml) are commonly studied in the literature in lung cells. While these findings provide insights into cellular responses, the overall environmental impact of PS-MNPs remains limited at realistic exposure levels.

**Highlights:**

- PS-MNPs are internalized into lung cells, with higher uptake in non-malignant cells.
- PS-NPs lead to increased migration, DNA damage, oxidative stress, and disturbed cell cycle progression.
- PS-NPs promote activation of survival pathways in non-malignant cells and lung organoids.
- Exposure of PS-NPs may cause potential implications for lung cancer development and progression.

## Introduction

Plastic products are widely distributed due to their tremendous utility in daily life. Reports indicate a steady annual increase in global plastic production, having reached nearly 400 million tons by 2023^1,2^. Inadequate recycling and the limited degradation of plastic materials pose significant environmental challenges. It is estimated that approximately 270 million tons of plastic waste are generated annually, with over eight million tons ending up in the ocean^3,4^. Under various aging conditions, such as biological degradation, physical-chemical abrasion, or ultraviolet radiation, plastic fragments degrade or break into smaller particles^5^. The precise distinction between micro-and nanoplastics (MNPs) remains debated. While some classifications define NP as particles smaller than 1 µm, stricter definitions specify particles below 100 nm^6,7^. The most commonly used polymers in these fragments include polyethylene (PE), polystyrene (PS), polypropylene (PP), polyethylene terephthalate (PET), and polyvinyl chloride (PVC), which are also prevalent constituents in the atmosphere^8^. Due to atmospheric transport, high MNP concentrations have been detected in numerous cities, remote areas such as mountain regions and the Arctic^9–12^.

Inhalation through the respiratory tract represents the primary route of MNP exposure compared to dermal contact or ingestion via contaminated food^13,14^. Elevated levels of airborne MNPs have been found in indoor and outdoor environments, originating from clothing fibers, automobile emissions, building materials, abrasive powders, and 3D printing^15–18^. However, limited sampling and methodological constraints have resulted in an incomplete understanding of airborne MNP abundance. Recent estimates suggest that individuals may inhale up to 70,000 airborne MNP particles each year, with fluctuations influenced by location, meteorological conditions, and occupation^19,20^.

Observational studies have identified a correlation between the accumulation of synthetic microplastic fibers and various lung dysfunctions among occupational workers^21,22^. Chronic exposure to manufactured MNPs has been linked to several respiratory diseases, including breathing difficulties, allergic alveolitis, and chronic pneumonia^23^. Due to their nanoscale dimensions, NPs exhibit an increased capacity for cellular penetration, allowing them to enter the bloodstream and accumulate in distant tissues, thereby raising concerns about long-term health issues^24,25^.

Lung cancer, the deadliest cancer worldwide, has been linked in some studies to exposure to specific types of NPs^26^. It is broadly categorized into two main subtypes: non-small cell lung cancer (NSCLC), which accounts for approximately 85% of cases, and small-cell lung cancer (SCLC), which comprises the remaining 15%^27,28^. Studies investigating the toxicity of PS-NPs with different sizes and surface charges in NSCLC cells (A549) have shown increased migration, upregulation of epithelial-to-mesenchymal transition (EMT) markers, and elevated reactive oxygen species (ROS) levels^29^. PS-NPs have been found to induce ferroptosis, disturb mitochondrial structure and function, and trigger autophagy, ultimately leading to lung injury in non-malignant lung epithelial cells^30^.

The aim of this study was to investigate the pulmonary toxicity of PS-MNPs in non-malignant and malignant human lung cells. We focused on assessing cellular uptake, migration, oxidative stress, and DNA damage to better understand the molecular mechanisms and potential risks of polystyrene particles. However, the precise mechanisms by which prolonged exposure to PS-MNPs contributes to lung disease development and progression as well as their potential carcinogenic effects remain poorly understood. While several studies have described the adverse effects of high MNP concentrations on non-malignant or malignant pulmonary epithelial and mesothelial cells^8,31,32^, the impact of lower, more physiologically relevant MNP concentrations and the direct comparison between non-malignant and malignant lung cells remain unexplored.

## Materials and methods

### Cell lines and lung organoids

Human bronchial epithelial cells (BEAS-2B), human non-small cell lung cancer (NSCLC; A549, H460) and human small cell lung cancer (SCLC; DMS53, H372) cell lines were purchased from the American Type Culture Collection (ATCC). All cell lines were cultivated in RPMI-1640 (with L-glutamine, Sigma, St. Louis, Missouri, USA) medium supplemented with 10% heat-inactivated fetal calf serum (FCS) and maintained in a humidified atmosphere with 5% CO_2_ at 37 °C. Trypsin-EDTA (0.25%), Mg^2^+ and Ca^2^+-free phosphate buffered saline (PBS), and dimethyl sulfoxide (DMSO) were obtained from Gibco (Grand Island, New York, USA). Analysis authenticated cells and checked for mycoplasma contamination (MycoAlert, Lonza, Basel, Switzerland).

The collection of patient tissue for the generation of lung organoids (LOs) was conducted in accordance with the guidelines approved by the Ethics Committee of the Medical University of Vienna (EK Nr: 2414/2020). Patient-derived LOs were previously established and characterized following the protocols outlined by Sachs et al.^33^. Briefly, after mechanical and enzymatic tissue distribution, epithelial cells were isolated and embedded in basement membrane extract (BME) under optimized pathway signaling. Fourteen days after splitting, organoids were released from BME by resuspending the BME domes in ice-cold advanced DMEM medium (Sigma). Organoids were re-seeded onto tissue culture plates without extracellular matrix support and were used for further experiments.

### Polystyrene micro-and nanoparticles

Commercially available types of unlabeled and fluorescent labeled polystyrene (PS) micro-and nanoparticles (MPs and NPs) with different sizes were purchased from MicroParticles GmbH (Berlin, Germany). PS-NPs with the diameter of 0.24 ± 0.01 μm (spherical, aqueous suspension at 2.5% w/v, green fluorophore-labeled ex/em = 502/518 nm, and aqueous suspension at 5% w/v, unlabeled) and PS-MP with the diameter of 1 µm (spherical, aqueous suspension at 2.5% w/v, red fluorophore labeled ex/em = 530/607 nm) were used in this study. The stock solutions were 25 mg/ml and 50 mg/ml in sterile water and were stored at 4 °C. For cell culture, working solution was freshly prepared in RPMI-1640 (Gibco, New York, USA) medium supplemented with L-glutamine, 10% FCS and vortexed for several minutes to avoid potential storage agglomerations.

### Cell viability assay (MTT assay)

Cells (5-10 x 10^3^) were seeded into 96-well culture plates and allowed to adhere overnight with 5% CO_2_ at 37 °C. PS-MNPs with ascending doses (0-60 µg/cm^2^) were prepared in fully supplemented RPMI-1640 culture medium and added on the following day. After 72 h, a tetrazolium-based colorimetric MTT assay (EZ4U, Biomedica, Austria) was performed to assess the cell viability as described^34^. Absorbance was measured at 450 nm and 620 nm using a Varioskan Lux plate reader (Thermo Fisher Scientific, Waltham, Massachusetts, USA).

### Colony formation assay

We performed colony formation assays to evaluate the effects of long-term exposure to PS-MNPs. Cells (2 x 10^3^) were seeded into 6-well culture plates and treated on the following day. After 7-14 days, cells were fixed with 70% ethanol for 30 min at RT and air dried. Colonies were stained with 0.01% (w/v) crystal violet for 1-2 h at RT. Excessive crystal violet was removed with distilled water. Microscopic images were taken with Evos M5000 (Thermo Fisher Scientific). For photometric measurements (562 nm), colonies were destained with a 2% SDS solution as described^35^. Triplicates were quantified on a Varioskan Lux plate reader (Thermo Fisher Scientific).

### Immunofluorescence

Cells (1-3 x 10^3^) were seeded into Ibitreat 8-well chamber slides (IBIDI, Gräfelfing, Germany) and exposed to two groups of plastics, with different sizes to fluorescent green and red labeled PS-MNPs (1 µg/cm^2^) the following day. After incubation for 24 h, cells were washed several times with 1x PBS to remove residual MNPs and fixed with 4% paraformaldehyde (PFA; Sigma-Aldrich, Germany) for 15 min. Cells were washed and blocked with 1% BSA in PBS for 30 min. Next, samples were incubated with the mouse monoclonal primary antibody targeting β-actin (13E5) rabbit mAB (Cell Signaling Technology; CST, #4970, 1:400 in PBS) for 1 h, followed by staining with secondary antibodies conjugated to Alexa Flour 488 (Thermo Fisher Scientific, #A-21206) for red PS-MP or Alexa Fluor 594 donkey anti-rabbit (#A-21207) for green PS-NP at a dilution of 1:1000 for 30 min at RT, respectively. 4′,6-diamidino-2-phenylindole (DAPI) was used at a dilution of 1:1000 in PBS to counterstain nuclei. Slides were mounted using Fluormount-G (Invitrogen, Thermo Fisher Scientific, Waltham, Massachusetts, USA) mounting medium. One day after being released from BME as described above, lung organoids were treated with labeled 1 or 10 µg/cm² PS-NPs for 14 days, untreated organoids serving as controls. Half of the medium was replaced every third day with fresh organoid medium containing plastic. Organoids were fixed in 4% PFA prepared in PBS for 30 min at RT on days 14 post-treatment. Fixed LOs were then transferred onto microscopy slides and stained with a 1× solution of CellMask™ Deep Red Actin Tracking Stain (A57245) for 30 min at RT. Nuclear staining and mounting were performed using VECTASHIELD® Antifade Mounting Medium with DAPI. Images were taken with a spinning microscope using a 63x oil immersion lens.

### Particle uptake and release

The internalization of PS-MNP particles was quantified by flow cytometry. Briefly, 24 h after seeding cells (5 x 10^5^) in 6-well culture plates, cells were exposed to three different doses of green PS-NP and red PS-MP (1, 10, and 25 µg/cm^2^) for further 24 h. After incubation, cells were washed twice with 1x PBS and collected in 200 µl FACS buffer (1x PBS + 2% FBS). For the release assay, cells were seeded as described above. Following a 24-hour exposure period with 10 µg/cm^2^ of each particle, the PS-MNPs-containing medium was replaced with fresh medium, and the remaining intracellular particles were quantified at various time points (2, 4, 6, 24, 48, and 72 h). DxFlex Flow Cytometer (Beckman Coulter, California, USA) was used for the analysis and approximately 10,000 events per sample were analyzed. The fluorescence’s fold change (FC) was normalized against the mean autofluorescence intensity of respective controls as a reference.

### Live-cell videomicroscopy, migration and cell fate analysis

Cells (2 x 10^3^) were seeded into 12-well plates and treated with 10 µg/cm^2^ unlabeled PS-NPs on the following day. Live-cell videos were generated using a Nikon Visitron Live Cell System (Visitron Systems GmbH, Puchheim, Germany) at 10 min intervals at a 10x magnification over 96 h. For cell fate maps, single cells (n = 25) were manually tracked using Image J, and the duration of Interphase and M-phase were calculated in Microsoft Excel. Cell migration and additional analyses of migratory behavior, including origin plots and nearest neighbor distance, were derived from the same live cell videos (single cells n > 280) using Image J and the Manual Tracking plugin.

### Apoptosis assay

Cells (3 x 10^5^) were grown in 6-well plates and treated on the next day with 10 µg/cm^2^ unlabeled PS-NPs. After various exposure times, cells were collected, washed with cold PBS, and resuspended in 100 µl 1x AnnexinV Binding Buffer containing 5 µl FITC-AnnexinV (556420, BD Pharmingen, California, USA) and propidium iodide (PI; HY-D0815, MedChemExpress). After incubation for 15 min at RT in the dark, staining was detected by flow cytometry (DxFlex, Beckerman Counter) at a wavelength of 488/530 nm for FITC-Annexin V and 535/617 nm for PI. At least 10,000 events per sample were counted and experiments were performed in three independent rounds.

### Cell cycle assay

Cells (3 x 10^5^) were seeded into 6-well plates and treated on the following day with PS particles with a concentration of 10 µg/cm^2^ at different time points. After washing with 1x PBS, cells were trypsinized, harvested, and centrifuged. Pellets were fixed with pre-cooled 70% ethanol (-20 °C) overnight at 4 °C. After centrifugation, cells were resuspended in 200 µl of PI solution (50 µg/ml, 500 µg/ml RNAse in PBS) and measured by flow cytometry (DxFlex, Beckerman Counter). Data were analyzed using the FlowJow v10.7.2 software (TreeStar, Ashland, Oregon, USA).

### TUNEL assay

For the detection of apoptotic cells undergoing extensive DNA degradation, a TUNEL reaction was carried out with an in-situ cell death detection fluorescein kit (Roche, Basel, Switzerland) according to the manufacturer’s instructions. Cells (1-3 x 10^3^) were seeded into Ibitreat 8-well chamber slides and treated the following day with 10 µg/cm^2^ unlabeled NP-PS for 24 h. Nuclei were counterstained with DAPI. Representative pictures were randomly taken on an inverse Micro Ti Eclipse FL microscope (Eclipse TI, Nikon). At least 150 cells per group were evaluated using Image J software and the percentage of TUNEL-positive cells per sample was assessed.

### Single-cell gel electrophoresis / comet assay

Cells (5 x 10^5^) were seeded in a 6-well culture plate and treated the following day with 10 µg/cm^2^ unlabeled PS-NP for 24 h. On the next day, cells were harvested and cell viability >70% was ensured using trypan blue. The standard comet assay for DNA damage detection was performed as described^36,37^. Briefly, collected cells were mixed with 0.5% LMPA (UltraPure Low Melting Point Agarose, Invitrogen) and transferred onto 1% normal agarose-coated slides. Following 1-hour lysis (2.5 M NaCl, 0.1 M Na2EDTA, 10 mmol/L Trizma base, pH 10, 1% Triton X-100), the process of DNA unwinding (30 min) and electrophoresis (30 min, 300 mA, 1.0 V/cm, at 4 °C) was conducted in an alkaline buffer (0.3 M NaOH, 1 mmol/L Na2EDTA, pH > 13). Subsequently, slides were washed 2x with dH_2_O (8 min), air-dried, and stained with PI (10 µg/mL). H_2_O_2_ (Sigma-Aldrich) at a concentration of 50 µmol/L was used as a positive control for 5 min on ice. Negative controls were treated with PBS. Cell analysis (150 nuclei) was evaluated under a fluorescence microscope (Nikon EFD-3) using a 10-fold objective. DNA migration was quantified using a computer-assisted image analysis system (Comet Assay IV, Perceptive Instruments). The percentage of DNA in tail (% DNA) was used as a parameter of DNA migration serving as the endpoint measurement.

### BER and NER repair enzyme (by use of modified comet assay)

A modified version of the comet assay was conducted to assess base excision repair (BER) and nucleotide excision repair (NER). These methods allow to detect the ability of repair proteins in cell extracts of different cell lines with and without PS-NP treatments to recognize and cleave substrate DNA containing specific lesions by use of different repair enzymes^38^.

Cells were plated as described above, and protein extracts were prepared by centrifuging samples at 700 g for 10 min at 4 °C after adding 200 µL of extraction buffer (45 mmol/L HEPES, 0.4 M KCl, 1 mmol/L EDTA, 0.1 mmol/L dithiothreitol, 10% glycerol, pH 7.8) with 1% Triton X-100, further described as buffer A. Samples were vigorously mixed, quickly frozen in liquid nitrogen, thawed, and subsequently centrifuged at 15,000 g for 5 min at 4 °C. 200 µl supernatant was collected and 60 µL of cold buffer B (40 mmol/L HEPES, 0.5 mmol/L EDTA, 0.2 mg/mL BSA, 0.1 M KCl, pH 8) was added. Protein concentrations were measured using the BCA Protein Assay Kit (Pierce). For BER measurements, the photosensitizer Ro 19-8023 (Chiron AS) which induces oxidation of DNA bases was used at 1.0 µmol/L. Substrate cells were treated in presence and absence of visible light (400 W, 60 cm distance, 4 min). For the NER assay, UVC (2.0 Jm−2, 22 s. on ice) was used to produce cyclobutane pyrimidine dimers. After light and UV treatment, cells were centrifuged (700 g for 10 min), and the resulting pellets were resuspended in freezing medium and cryopreserved at-80°C. A549 cells were used as substrate cells and were treated for the assay as previously described^39^. Pre-treated substrate cells (2 x 10^4^ per gel) were embedded in agarose and lysed for 1 h. For NER measurements, slides were washed twice for 10 min in buffer N (45 mmol/L HEPES, 0.25 mmol/L EDTA, 0.3 mg/mL BSA, 2% glycerol, pH 7.8), and buffer B was used for BER. Nuclei were then incubated for 30 min with either 50 µl of “extract mix” or 50 µl of control buffer. For BER experiments, formamidopyrimidine-DNA glycosylase (FPG) was used as the positive control, and T4EndoV served as the positive control for NER measurements. Negative controls contained buffer B for BER and buffer N for NER. Alkaline unwinding and electrophoresis were conducted following standard comet assay protocols as described above.

### ROS and GSH/GSSG detection

The reactive oxygen species (ROS) assay was performed using a commercially available ROS-Glo^TM^ H_2_O_2_ assay kit, according to the manufacturer’s instructions (Promega, Madison, Wisconsin, USA). Briefly, cells (5-10 x 10^3^) were seeded onto opaque white 96-well plates and were exposed to two different concentrations of unlabeled PS-NPs (1 and 10 µg/cm^2^) for 18 h on the following day. H_2_O_2_ substrate solution was then added to achieve a final volume to 100 µl, and plates were incubated for an additional 6 h. After 24-hours of treatment, 100 µl of the ROS-Glo detection solution was added to each well and incubated at RT before measurement for 20 min. The glutathione ratios were analyzed with the GSH/GSSG-Glo^TM^ assay (Promega). Cells (5-10 x 10^3^) were seeded and treated on the next day with 10 µg/cm^2^ unlabeled PS-NP for 24 h. The total glutathione (GSH + GSSG) and the oxidized form (GSSG) were quantified in two parallel reactions. In both assays, the luminescent light signal produced was proportional to the levels of H_2_O_2_ and GSH in the cells. Luminescence was recorded with the Varioskan (Thermo Fisher Scientific).

### Quantification of lysosomes

Cells were grown in 12-well plates and then incubated with 1 µg/cm^2^ green labeled PS-NP for 24 h. After washing and harvesting, cells were stained with LysoTracker® Deep Red(Invitrogen, Thermo Fisher Scientific) for 30 min and DAPI. The localization of lysosomes and labeled PS-NPs was evaluated using Amnis® imaging flow cytometry (Merck, Darmstadt, Germany). A total of 10,000 events per sample was counted, all experiments were performed in two independent rounds.

### Protein isolation and western blots

Cells (5 x 10^5^) were seeded into 6-well plates and collected after 24 h of incubation with 10 µg/cm^2^ unlabeled PS-NPs following harvesting in RIPA buffer supplemented with Protease Phosphatase Inhibitor Cocktail (Thermo Fisher Scientific). LOs were exposed to 50 µg/cm^2^ unlabeled PS-NPs for 14 days before they were dissociated and harvested in RIPA. Protein concentrations were determined using the Pierce BCA assay (Thermo Fisher Scientific). Protein extractions (10 µg) were separated using Mini-PROTEAN Tetra electrophoresis cells (Bio-Rad Laboratories Hercules, California, USA) and subsequently transferred to a nitrocellulose membrane (Amersham Protran, GE Healthcare) by Trans-BlotTurbo (Bio-Rad Laboratories). Membranes were blocked with 5% non-fat dry milk in 1x TBS-T (Tween 0.1%) for 1 h. Immunodetection was performed using Super Signal West Femto Chemiluminescent Substrate (Thermo Fisher Scientific). The following primary antibodies were used at a concentration of 1:1000 overnight at 4 °C: pERK1/2 (Thr202/Tyr204, D13.14.4E, #4370, CST), ERK1/2 (137F5, #4695, CST), pAKT (Ser473, D9E, #4060, CST), and AKT (C67E7, #4691, CST). GAPDH (#5174, CST, 1:10000) was used as a loading control and secondary antibodies were purchased from Dako and incubated at a dilution of 1:1000 for 2 h at RT. Chemiluminescence imaging was conducted on a Vilber Fusion FX system. Band intensity quantification was performed with Image J.

### Phosphorylation antibody array for receptor tyrosine kinases (RTKs)

BEAS-2B cells (5 x 10^5^) were seeded in 6-well plates and treated with 10 µg/cm^2^ unlabeled PS-NPs for 24 h. Cells were collected and measured as described above. The human RTK Phosphorylation Antibody Array Membrane (ab193662, Abcam, Cambridge, UK) was used to determine the phosphorylation of 71 RTKs according to the manufacturer’s instructions. In brief, antibody array membranes were blocked and cell lysates were incubated overnight at 4 °C under gentle agitation. Membranes were incubated with biotinylated anti-phosphotyrosine kinases following HRP-conjugated streptavidin for 2 h at RT. Membranes were washed thoroughly between each incubation step. Spot signals were quantified using Image J and normalized to control spots.

### Statistical analysis

Unless stated otherwise, each experiment was repeated three times, and values are given as mean ± standard error of the mean (SEM) of three independent biological repeats performed in technical triplicates, which are normalized to the untreated control, respectively. Differences were analyzed by Student’s t-Test or ANOVA including Dunńs or Tukey’s multiple comparisons test in GraphPad Prism 8.0. The p-value ≤ 0.05 was considered statistically significant.

## Results

### PS-MNPs did not induce cell death in non-malignant and malignant lung cell lines

To assess the potential toxicity of PS particles in our cell models, we exposed cell lines to increasing concentrations of two PS-MNPs which were commonly studied in the literature (0, 15, 30, and 60 µg/cm^2^, equals to 0, 25, 50, and 100 µg/ml) and tested cell viability after 72 h using MTT-based assays. In this study, we refer to microplastics (MPs, 1 µm – 5 mm) and nanoplastics (NPs, < 1 µm diameter). Reduced cell viability by a maximum of 20% was observed only at very high doses (> 30 µg/cm^2^) of PS-MNP treatment, with the non-malignant BEAS-2B cells showing the strongest dose-dependent response (Fig. 1A). Similar results were obtained using unlabeled PS-NPs by MTT-based and SYBR green-based growth assays (Supplementary Fig. S1). To stay as close as possible to low physiological conditions, we selected 1 and 10 µg/cm^2^ concentrations (equal to 1.65 and 16.5 µg/ml), for further experiments (indicated by the arrows), as they did not cause any growth inhibition. Colony formation assays showed no significant effects on proliferation or growth after long-term exposure (10 days) to 10 µg/cm^2^ PS-MNPs (Fig. 1B, Supplementary Fig. S2).

**Figure 1:**
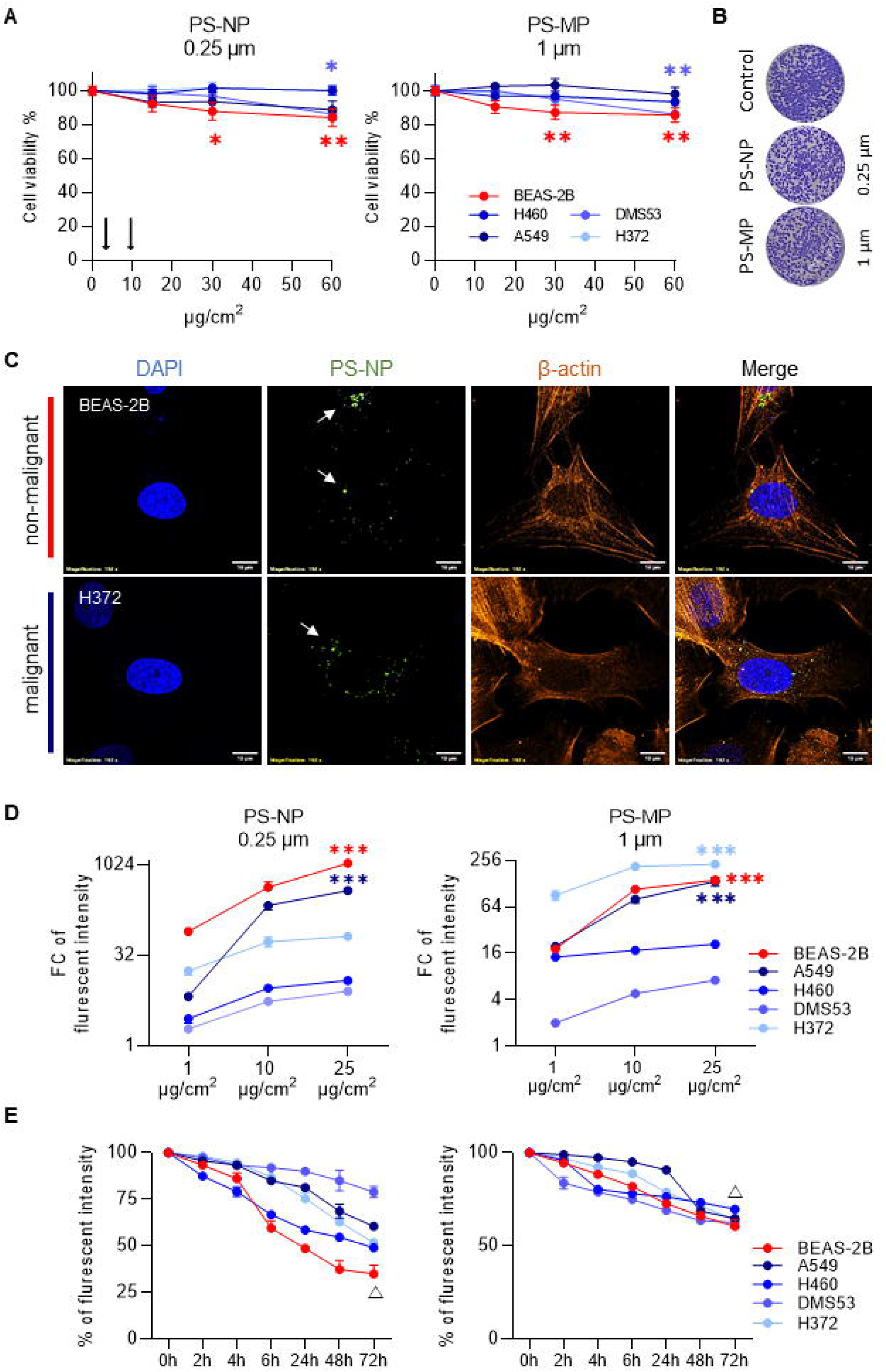
**PS-MNP induced no strong impact on cell viability but higher levels of uptake and release in non-malignant (BEAS-2B) lung cells**. **A)** Cell viability was assessed using an MTT-based assay after 72 h of exposure to increasing doses (0-60 µg/cm^2^) of PS-MNPs of indicated sizes (0.25 µm / 1 µm). ANOVA and Dunnett’s multiple comparisons test. *p ≤ 0.05 and **p ≤ 0.01. B) Representative pictures of colony formation assays for BEAS-2B cells after a 10-day incubation with 10 µg/cm² of both PS-MNPs. C) Confocal microscopy images showed the cellular localization of green fluorescent PS-NPs (arrows) at 1 µg/cm^2^ concentration after 24 h. Non-malignant (BEAS-2B) and malignant (H372) cells were used. The cytoskeleton was stained with β-actin (red) and DAPI (blue), indicating the cell nucleus. Scale bar: 10 µm. D) The uptake of both PS-MNPs was assessed using flow cytometry. The fold change (FC) of fluorescence was normalized relative to the average fluorescence of control, which served as a reference. ANOVA and Tukey’s multiple test compared to 1 µg/cm². ***p ≤ 0.001. E) Quantitative fluorescence analysis of PS-MPS release. The % of fluorescent intensity represents the number of respective particles retained within cells. Each experiment was repeated three times, and data are shown as mean ± SEM compared with respective untreated (0 h) control cells. ANOVA and Dunnett’s multiple comparisons test. The triangle (,6) indicates ****p ≤ 0.0001 for all cell lines.

### Non-malignant lung cells exhibit the highest PS-NP accumulation and release

Nevertheless, we detected numerous PS-MNP particles inside the cells, predominantly accumulating in the cytoplasm, even at the low concentration of 1 µg/cm^2^ (Fig. 1C, Supplementary Figs. S3A and S3B). Flow cytometry analysis of cellular uptake kinetics demonstrated that both PS particle sizes were internalized in a dose-dependent manner after 24 h of exposure (Fig. 1D). Notably, a significantly stronger signal of 0.25 µm PS-NP was observed in non-malignant BEAS-2B cells compared to other cell lines, suggesting that BEAS-2B have a greater tendency to accumulate smaller PS particles.

To better understand particle dynamics, we investigated whether cells can release internalized PS-MNPs (Fig. 1E). In all cell lines, intracellular levels of 0.25 µm PS-NPs (right) declined after 6 h, reaching a plateau after 24 h. After 72 h, approximately 55% of PS-NPs remained in malignant cells, while non-malignant cells retained about 35%. A similar pattern was observed for 1 µm PS-MPs (left), with around 60% of PS-MPs remaining after 72 h. These observations indicate that the cells neither degraded nor expelled a substantial proportion of internalized PS-MNPs. Only the smallest PS-NP showed the most pronounced decrease in non-malignant BEAS-2B cells. Based on these results, further assays will focus on a more detailed examination of the effects of small PS-NP particles.

### PS-NPs induced cell migration, scattering, and S-phase arrest

To further elucidate the potential impact of PS particles, we analyzed the migratory behavior using live-cell video microscopy and single-cell tracking over 96 h. All cell lines, except one (H460), exposed to PS-NPs displayed significantly increased total migrated distance compared to untreated controls, which is also reflected in the origin plots (Figs. 2A, 2B, and Supplementary Fig. S4). Interestingly, cells also showed a significant increase in cell scattering after 24 h of PS-NP treatment, quantified by nearest neighbor distance (Fig. 2C). Since increased migration and scattering are associated with pathological effects, we further analyzed the cell cycle distribution. We found that 10 µg/cm^2^ PS-NP effectively inhibited cell cycle progression in our cell lines. After 72 h of exposure, a significant time-dependent increase in S-phase and G2/M phase was observed in both non-malignant and most malignant cell models. In contrast, no significant changes were noted in A549 (Figs. 2D and 2E).

**Figure 2:**
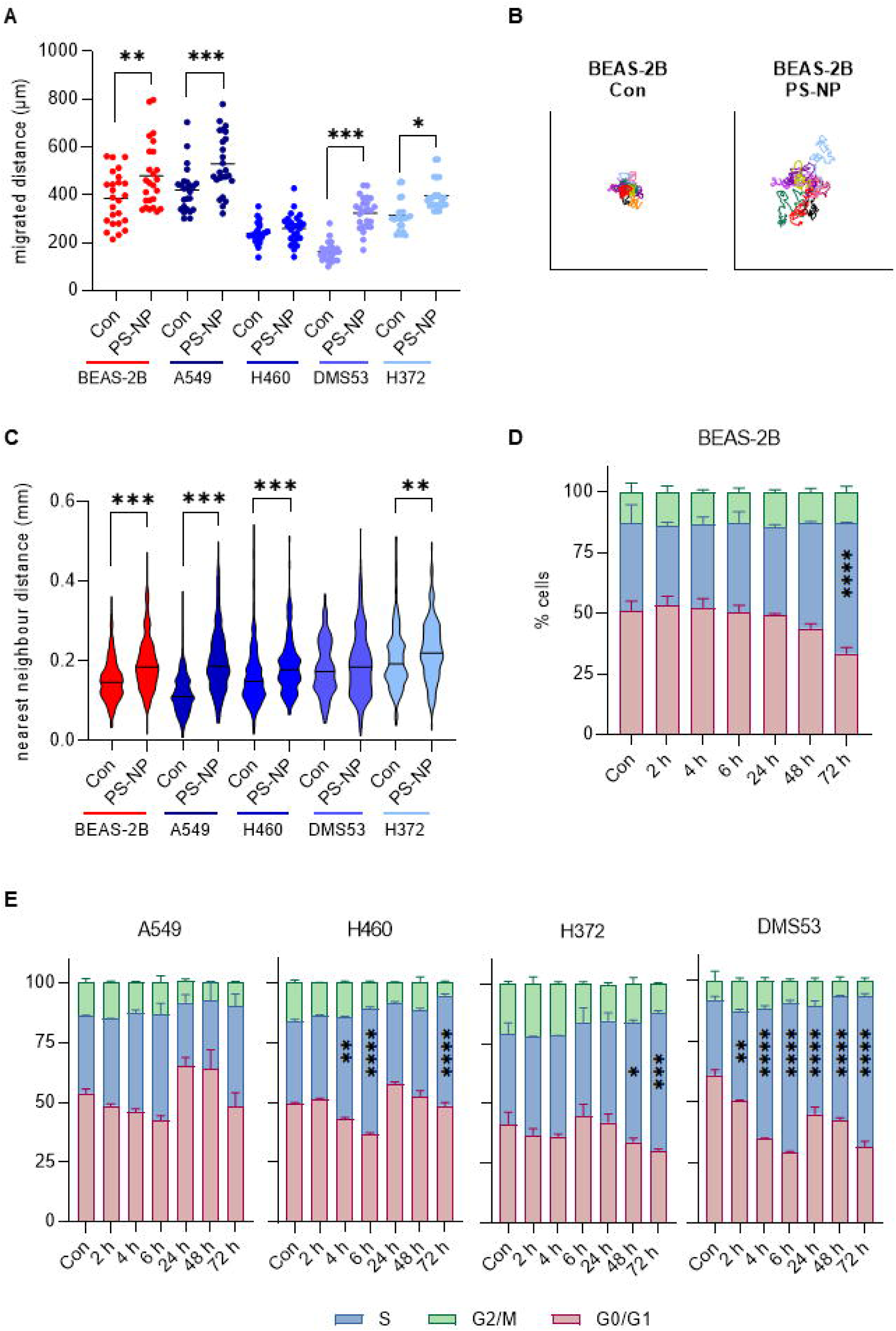
PS-NPs induced cell migration, scattering, and S phase arrest. **A)** Migrated distance of all single cells over 96 h of exposure to 10 µg/cm² unlabeled PS-NP, quantified by manual single cell tracking in Image J. Each dot represents an individual cell, with the mean indicated by the horizontal lines. Ordinary One-Way ANOVA. **B)** Origin plots of representative tracked BEAS-2B cells (n = 10) were calculated by Image J. **C)** Distance to the nearest neighbor of > 280 single cells derived from live cell imaging after 24 h. Data are shown as mean ± SEM. Kruskal-Wallis test. **D)** and **E)** Cell cycle distribution was analyzed via flow cytometry after different time points of unlabeled 10 µg/cm² PS-NP treatment in D) non-malignant and E) malignant cell lines. Experiments were performed in triplicate and bars show mean ± SEM compared with respective untreated control cells (red - G0/G1, blue - S, green - G2/M). ANOVA followed by Dunnett’s multiple comparisons test. *p ≤ 0.05, **p ≤ 0.01, ***p ≤ 0.001, and ****p ≥ 0.0001.

For a deeper characterization, we generated cell fate maps of individual cells using previously generated videos (Supplementary Fig. S5). Comparing untreated control cells with those treated with 10 µg/cm² PS-NPs, we observed no significant changes indicative of abnormal cell proliferation, which is consistent with our cell proliferation data in Fig 1. Flow cytometry-based apoptosis assays using Annexin V and PI as markers for early, late apoptotic, and dead cells at various time points confirmed that 10 µg/cm^2^ PS-NPs have no significant impact on cell death (Supplementary Fig. S6).

### Impact of PS-NPs on DNA damage and on repair capacity in malignant and non-malignant lung cell lines

Since S-phase arrest is a strong indicator of DNA damage or disruption, we conducted TUNEL staining to detect DNA double-strand breaks. Cells were exposed to 10 µg/cm² PS-NPs for 24 h, fixed, and stained for TUNEL reaction. Our data revealed significantly increased DNA damage in PS-NP-treated cells, which aligns with the observed prolonged S-phases in these cells (Fig. 3A).

**Figure 3:**
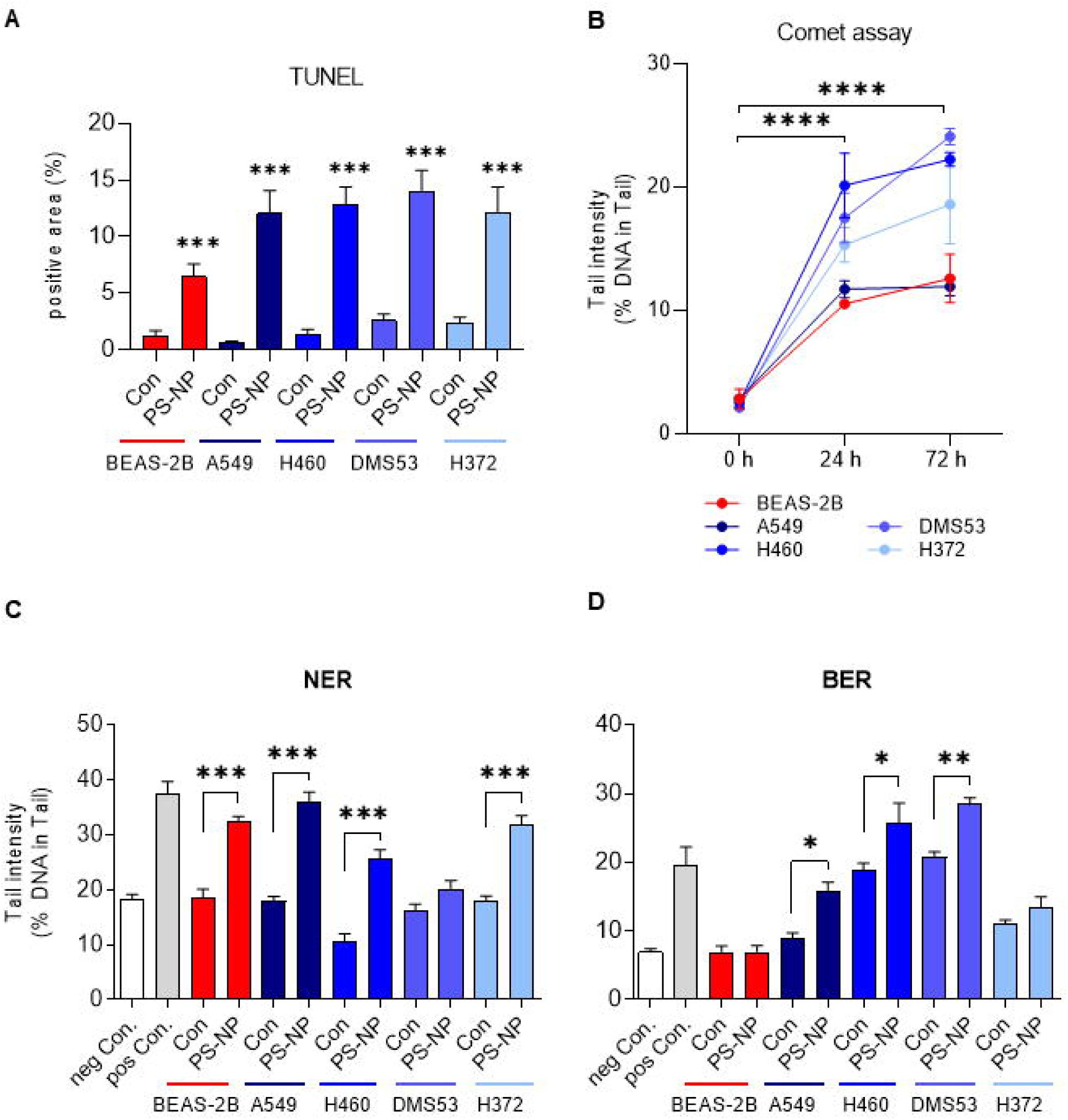
Enhanced DNA damage and altered DNA repair capacity induced by PS-NP treatment. **A)** Quantifying TUNEL-positive cells after 24 h of exposure to 10 µg/cm² PS-NP (green). Between 6 and 15 representative images were evaluated using Image J. ANOVA and Sidak’s multiple comparisons test. **B)** DNA damage was measured in all lung cells using the standard comet assay. Determined baseline DNA damage levels after 24 and 72 h treatment with 10 µg/cm² unlabeled PS-NP are expressed as a percentage of tail intensity relative to untreated cells. H_2_O_2_ (50 µmol/L) was used as a positive control. ANOVA and Dunnett’s multiple comparisons test. ****p ≥ 0.0001. **C)** and **D)** Repair enzyme experiments with all lung cell lines elucidating the NER and BER capacity compared to untreated control cells, respectively. FPG for BER and T4EndoV for NER were used as positive controls. ANOVA and Sidak’s multiple comparisons test. *p ≤ 0.05, **p ≤ 0.01, and ***p ≤ 0.001. Data are shown as mean ± SEM of 6 microscopic fields of view from at least two independent experiments.

Comet assays reflecting single-strand, double-strand, and apurinic site breaks confirmed DNA damage^40^, as indicated by significantly higher tail intensities in PS-NP (10 µg/cm²) exposed cells after 24 and 72 h (Fig. 3B, Supplementary Fig. S7). We performed a modified comet assay to measure nucleotide excision repair (NER) and base excision repair (BER) capacity.

Our findings revealed that PS-NP treatment led to increased NER activities in non-malignant and some tumor cells (Figs. 3C). In non-malignant BEAS-2B cells, NER was significantly elevated after PS-NP treatment, whereas BER levels remained unchanged in contrast to the increased BER levels observed in some malignant cells (Fig. 3D).

### PS-NPs caused oxidative stress and triggered the antioxidant system in non-malignant lung cells

Since DNA damage is often associated with oxidative stress, we measured reactive oxygen species (ROS) production. We observed an increase of ROS levels in several cell lines (BEAS-2B, H460, and H372) after 24 h of PS-NP treatment (Fig. 4A). In order to understand the adaptive response, we quantified the GSH/GSSG ratio by comparing the glutathione levels (µM) without and with 24 h of PS-NP exposure (10 µg/cm^2^). Significantly, the oxidative stress response and antioxidant capacity only dramatically changed in the non-malignant lung cell line after PS-NP treatment (Fig. 4B). Total glutathione (GSH) and oxidized glutathione (GSSG) levels were higher in malignant lung cells at baseline compared to the non-malignant cell, reflecting the cellular stress response of tumor cells (Supplementary Fig. S8). Regarding intracellular particle localization, our imaging data from AMNIS flow cytometry revealed a co-localization of lysosomes (red) with PS-NPs (green) in all cells, indicating that PS-NPs are taken up by lysosomes (Fig. 4C, Supplementary Fig. S9). However, no significant alterations in lysosomal numbers were observed (Supplementary Fig. S10).

**Figure 4:**
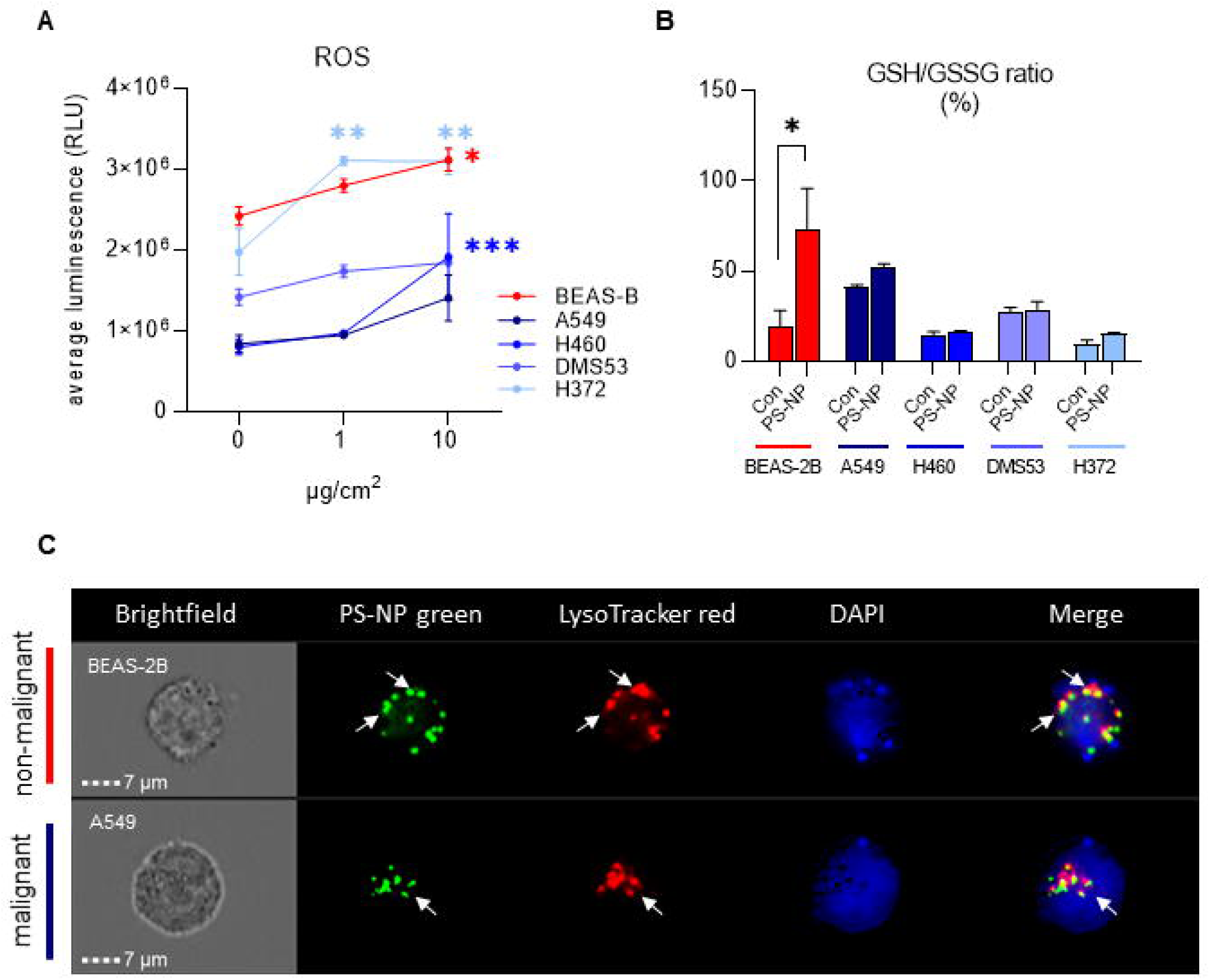
Increased ROS and impaired intracellular redox balance in lung cells upon PS-NP. **A)** ROS production in all lung cells after 24 h of incubation with 1 and 10 µg/cm² unlabeled PS-NP was measured using a luminescence-based assay (Promega). **B)** The GSH/GSSG ratio (%) of all cell lines was determined using a luminescence-based GSH/GSSG-Glo assay (Promega). Cells were treated with PS-NP (10 µg/cm², 24 h) and data are shown as the mean ± SEM of at least two independent experiments. ANOVA and Tukey’s multiple comparisons test. *p ≤ 0.05, **p ≤ 0.01, and ***p ≤ 0.001 compared to non-treated control cells, respectively. **C)** Representative images of non-malignant (BEAS-2B) and malignant (A549) lung cells incubated with PS-NP (green, 10 µg/cm², 24 h), LysoTracker (red), and DAPI (blue), acquired using AMNIS flow cytometry. Scale bar: 7 µm.

### Increased phosphorylation of AKT and ERK triggered by PS-NP exposure in non-malignant cells

Encouraged by our findings that the effects of PS-NPs were most pronounced in non-malignant cells and supported by evidence showing that these cells exhibit an increased response to oxidative stress, we investigated the activation of major survival and mitogenic pathways. Our results revealed distinct phosphorylation patterns between non-malignant and malignant cells following 24 h of treatment with 10 µg/cm² PS-NPs (Fig. 5A). Interestingly, non-malignant BEAS-2B cells exhibited dramatically increased levels of pAKT and pERK which indicates the activation of survival mechanisms or a stress response triggered by PS-NPs. In contrast, tumor cells showed no changes in AKT or ERK phosphorylation, supporting our hypothesis that non-malignant lung cell lines may be more susceptible to PS-NP induced alterations (Fig. 5A and 5B).

**Figure 5:**
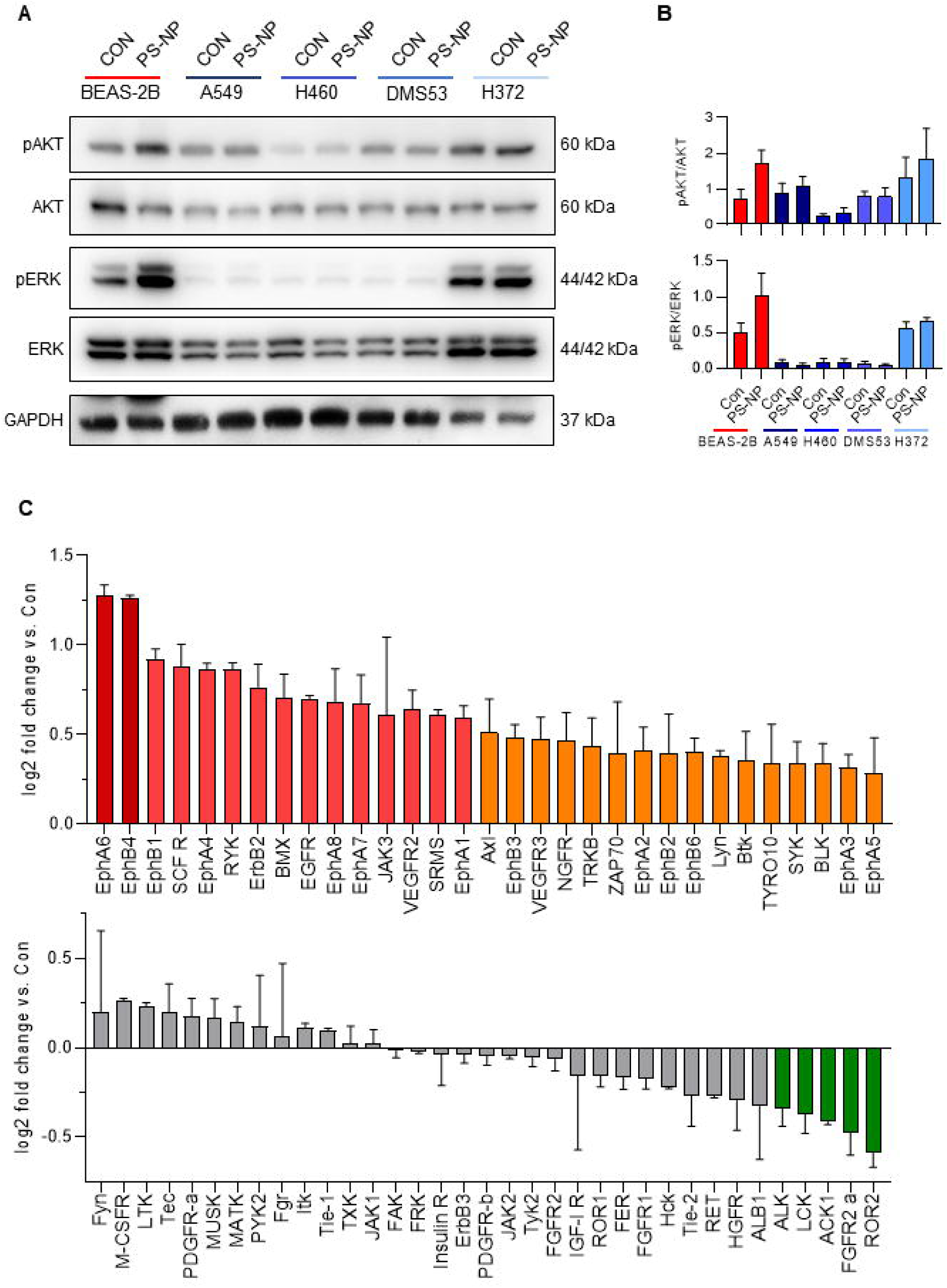
Dysregulated phosphorylation of AKT and ERK pathways following PS-NP treatment. **A)** Representative immunoblots of non-malignant and malignant lung cell lines treated with PS-NP (unlabeled, 10 µg/cm², 24 h), showing phosphorylated AKT and ERK relative to total AKT and ERK levels. **B)** Densitometric quantification was performed using Image J of at least three repeats. Phosphorylated proteins were normalized to total proteins, and the results shown are mean ± SEM. GAPDH was used as a loading control. **C)** Results of RTK assay in BEAS-2B cells treated with 10 µg/cm^2^ PS-NP for 24 h. The mean log2 FC was calculated from the mean pixel intensities compared to the untreated control cell.

To further explore the changes in the non-malignant BEAS-2B cells, we performed a human receptor tyrosine kinases (RTK) array. Comparison of BEAS-2B cell lysates without and with 24 h PS-NP treatment at 10 µg/cm^2^ revealed increased activation of ephrin (Eph) receptors (EphA6, EphB4, and EphB1), human epidermal growth factor receptor (HER2; ErbB2), stem cell factor receptor (SCFR), epidermal growth factor receptor (EGFR), vascular endothelial growth factor receptor (VEGFR2), JAK3, and AXL, potentially contributing to the activation of oncogenic pathways such as AKT and ERK (Fig. 5C, Supplementary Fig. S11).

### PS-NPs activated signaling pathways in 3D lung organoid models

Since our cell culture data showed that PS-NP treatment induces significant cellular changes predominantly in non-malignant human lung cells, we sought to validate these findings in a more physiologically relevant system. We used patient-derived lung organoids (LOs) to investigate the internalization of PS particle in a three-dimensional model. The LOs were generated following established protocols and consist of a polarized, pseudostratified epithelium comprising basal, secretory, and multi-ciliated cells. Indeed, after 14 days, PS-NPs were detected within the epithelial cell layers at both concentrations of 1 µg/cm² (Fig. 6A) and 10 µg/cm² (Fig. 6B) and in multiple focal planes, confirming the ability of particles to penetrate multiple cellular barriers. Furthermore, phosphorylation of both AKT and ERK was markedly elevated by PS-NP treatment in LOs, consistent with our previous findings in non-malignant BEAS-2B cells (Fig. 6C). Taken together, our data indicate that PS-NP exposure influenced crucial cellular response activity, potentially contributing to malignant transformation.

**Figure 6:**
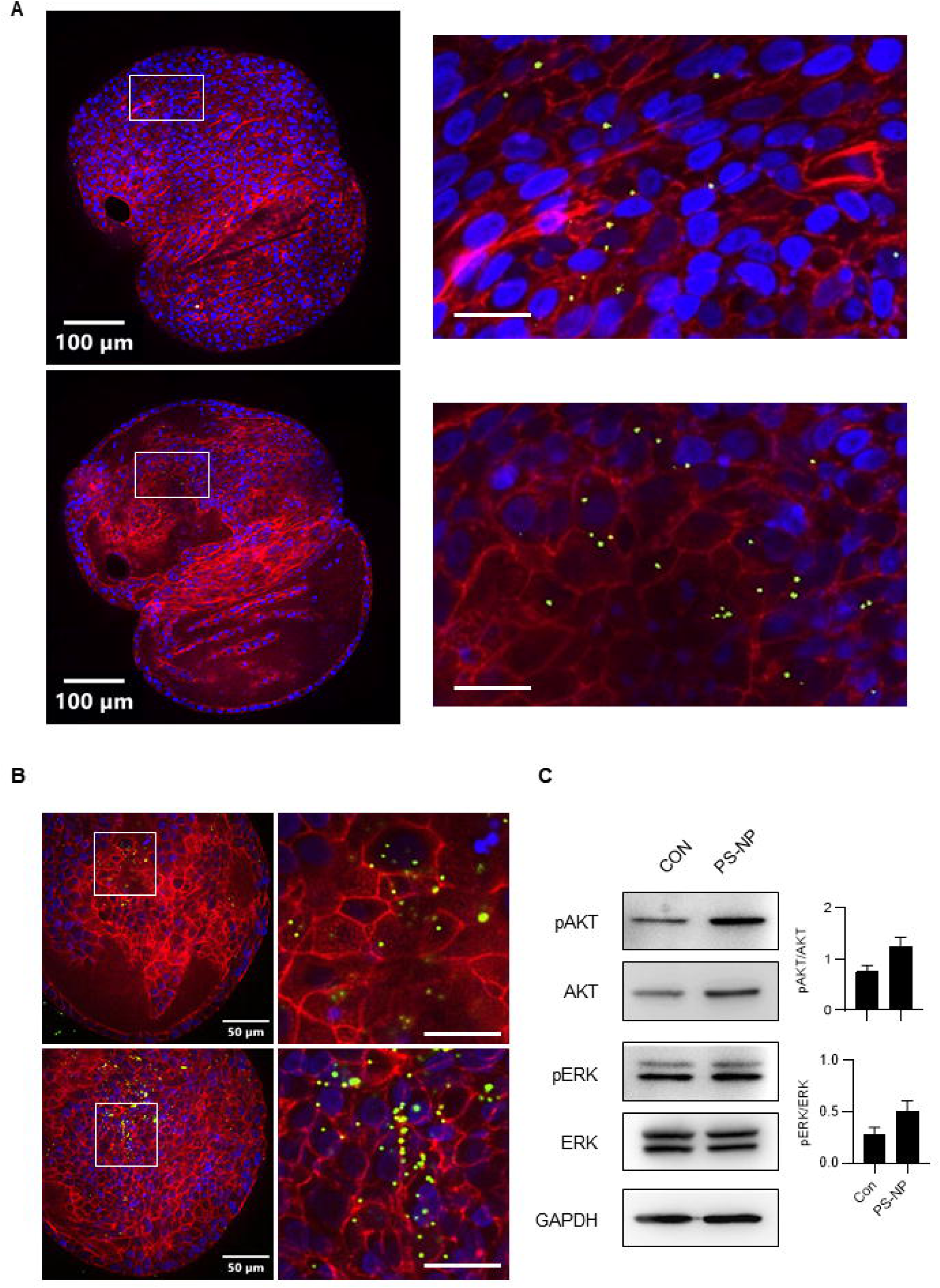
PS-NP uptake confirmed in lung organoids (LOs) and increased pAKT and pERK. Representative confocal images of LOs after 14 days incubation with **A)** 1 µg/cm² and **B)** 10 µg/cm² fluorescent green labeled PS-NP. Cells (actin) are shown in red, and nuclei (DAPI) in blue. Scale bars: A: 100 µm (left) and 20 µm (right); B: 50 µm (left) and 20 µm (right). **C)** Representative images from western blotting of LOs untreated or treated with PS-NP (50 µg/cm²) for 14 days. Protein expression of phosphorylated AKT and ERK relative to total AKT and ERK levels were measured. Protein densities were derived from at least two repeats. Phosphorylated proteins were normalized to total proteins, and the results shown are the mean ± SEM. GAPDH was used as a loading control.

## Discussion

In this study, we examined the effects of polystyrene micro-and nanoplastics (PS-MNPs) on non-malignant (BEAS-2B) and malignant (A549, H460, DMS53, and H372) lung cells, as well as human-derived lung organoids (LOs), whereas we focused on cell viability, internalization, migration, DNA damage, oxidative stress, and survival signaling pathways. Our findings reveal a complex interplay between PS-MNP exposure and cellular responses that differ significantly between non-malignant and malignant lung cell lines.

We found that PS-MNPs did not induce significant cytotoxicity in human lung cells at low concentrations. In this study, exposure to PS-MNPs below 30 µg/cm^2^ (equivalent to 50 µg/ml) had no significant effect on cell viability. In contrast, higher concentrations caused a modest cytotoxic effects which is consistent with previous reports^8,24^. Notably, non-malignant BEAS-2B cells exhibited the most pronounced and significant dose-dependent responses, highlighting their sensitivity to PS-MNP exposure^31,41,42^. Supporting these findings, a recent study using μFTIR spectroscopy identified MPs in 11 of 13 human lung tissue samples, with an adjusted mean concentration of 0.69 ± 0.84 MP/g. The dominant polymers, polypropylene (PP) and polyethylene terephthalate (PET) accounted for 23% and 18% of MPs, respectively. MPs were present across all lung regions, with the highest levels in the lower lung, highlighting inhalation as a likely exposure route^43^. In this context, we selected two relatively low concentrations with no cytotoxic activity for our subsequent analyses to stimulate a more realistic exposure scenario and investigate cellular alterations induced by PS-MNPs.

In our study, no significant inhibitory effects on cell growth following long-term treatment to 10 µg/cm^2^ PS-MNP over 10 days were observed. Previous in vitro studies have suggested that prolonged exposure to higher MNP concentrations (≥ 25 µg/ml) may better simulate chronic environmental exposure. Indeed, six-month PS-NP exposure studies have demonstrated early and adverse tumor-associated phenotypes, indicating that much longer exposure time may be required to reveal such effects^44,45^.

Despite limited cytotoxicity, PS-MNPs were efficiently internalized by both non-malignant and malignant lung cells in a dose-dependent manner within 24 h, predominantly accumulating in the cytoplasm. Notably, non-malignant BEAS-2B cells exhibited the highest internalization rates for smaller PS-NPs, underscoring their vulnerability to smaller plastic particles. Furthermore, kinetic assays demonstrated that 0.25 µm PS-NPs were accumulated and released more rapidly than 1 µm PS-MPs when cells were exposed to equal amounts of two different particle sizes, aligning with previous findings that smaller particles are more conductive to uptake^24,29^.

Our findings revealed that PS-NPs treated cells showed enhanced cell migration and more significant scattering behaviors in nearly all cell lines, which is in accordance with other studies. Recent reports have shown that exposure of different concentrations of PS-NP cause a significant increase of migratory activities in human alveolar epithelial and colorectal cancer cells^29,46^. Cell migration is a critical initiating event in the formation of metastases, making it a key hallmark of cancer aggressiveness and disease progression^47,48^. Additionally, PS-NP exposure has been shown to alter adhesion molecule expression, disrupt the extracellular matrix, and promote epithelial-to-mesenchymal transition (EMT)-like behavior^29,44,49,50^. These effects suggest that PS-NPs can disrupt normal cellular processes, potentially promoting features commonly associated with tumor aggressiveness and metastasis^44,51^.

We observed that PS-NP treatment at 10 µg/cm² concentration induced a significant, time-dependent increase of S-phase populations in several cells, indicating cell cycle arrest. The cell cycle is tightly regulated to ensure proper DNA replication, repair, and division. Disruptions in this process can lead to severe consequences, ranging from impaired cellular function to the onset of malignancy^52^. This observation aligns with previous *in vitro* studies and functional enrichment analyses demonstrating that several proteins involved in the cell cycle are upregulated following PS-NP treatment, resulting in altered cell cycle distributions^24,53,54^. Regarding the possible genotoxic effects on the molecular levels from PS-NPs exposure, our data show that PS-NPs induce DNA damage in all tested lung cell lines. These findings are consistent with our observations of dysregulated cell cycle progression. Observed S-phase arrest suggests DNA-replication stress or damage, while accumulation in the G2/M phase indicates an activation of checkpoint response to genomic instability^55^.

DNA damage and cell cycle disruption are frequently linked to oxidative stress. Indeed, PS-NP treatment led to a dose-dependent increase in reactive oxygen species (ROS) production in our cell lines. Excessive ROS levels are the products of high-stress reactions that contribute to DNA damage^56,57^. The antioxidant system is activated as a compensatory response. When antioxidant enzymes cannot counterbalance the large amounts of ROS, an imbalance between ROS generation and the antioxidant defense system occurs, consequently leading to increased oxidative stress^58^. Several earlier reports have identified oxidative stress as a central mechanism underlying the effects of PS particles in lung and intestinal cells^25,32,42,50^. Furthermore, oxidative stress has been linked to lysosomal damage and dysfunctions^41^, although our study did not observe significant changes in the lysosome amount after PS-NP treatment. Nevertheless, our results demonstrated that PS-NPs accumulated in the lysosomes of all lung cells after 24 h of exposure.

Interestingly, we observed dramatic shifts in antioxidant capacity only in non-malignant BEAS-2B lung cells. Changes in the GSH/GSSG ratios indicate a shift in the redox balance, which is crucial for cell survival and function. While earlier studies have shown that MNP exposure leads to decreased levels of antioxidant enzymes such as SOD, GSH, and AOC^50,51,53,59^, the prominent antioxidant response seen in our study suggests that BEAS-2B cells have an enhanced activation of compensatory mechanisms to counteract oxidative stress. This supports our hypothesis that normal cells experience significant cellular stress induced by PS-NPs. Although non-malignant cells are generally less stressed under normal conditions (with lower baseline total glutathione levels) compared to malignant cells, they appear more sensitive (with a higher GSH/GSSG ratio) to external stressors such as PS-NP exposure.

Our findings also indicated that PS-NP affects DNA repair capacity. Malignant cells demonstrated elevated base excision repair (BER) activities, likely an adaptive response to DNA damage. In contrast, non-malignant BEAS-2B cells exhibited insufficient BER activity, underlining the described vulnerability to genomic instability under PS-NP exposure in bronchial epithelial cells of Jean et al. 2023^41^. Impaired BER capacity, coupled with an increased GSH/GSSG ratio suggests that non-malignant lung cells may struggle to cope with PS-NP-induced genotoxic stress, which could affect their function and viability.

Finally, we observed that phosphorylation of AKT and ERK was elevated exclusively in non-malignant (BEAS-2B) cells. This activation of key survival and mitogenic signaling pathways suggests that PS-NP exposure may trigger survival mechanisms^31,60^. Furthermore, the receptor tyrosine kinase (RTK) array indicated increased activations of specific receptors, including Eph, EGFR, and VEGFR2 as mediators of this response. These pathways are critically involved in regulating proliferation, survival, and differentiation, indicating that PS-NPs may create a cellular environment conducive to tumor progression, particularly in non-malignant lung cells. Supporting these findings, our patient-derived lung organoid (LO) models demonstrated the ability of PS particles to translocate through cellular barriers in a three-dimensional setting and the activation of survival pathways as evidenced by increased pAKT and pERK levels.

Collectively, our results suggest that PS-NPs may not have significant effects on cancer cells but could potentially exhibit tumor-promoting effects on non-malignant cells by altering cellular stress response and survival mechanisms. Importantly, this study is the first to directly compare the impacts of PS particles on malignant and non-malignant lung cells, addressing a critical gap in the current understanding of the impact of plastic particles on lung cancer development and progression.

## Abbreviations

BER: Base excision repair
BME: Basement membrane extract
EMT: Epithelial-mesenchymal transition
ER: Endoplasmic reticulum
GSH: total glutathione
GSSG: oxidized glutathione
Los: Lung organoids
MNPs: Micro-and nanoplastics
NER: Nucleotide excision repair
NSCLC: Non-small cell lung cancer
PE: Polyethylene
PET: Polyethylene terephthalate
PP: Polypropylene
PS: Polystyrene
PS-MNPs: Polystyrene-micro-and nanoplastics
PS-MPs: Polystyrene-microplastics
PS-NPs: Polystyrene-nanoplastics
PVC: Polyvinyl chloride
ROS: Reactive oxygen species
RT: Room temperature
SCLC: Small cell lung cancer

## Conclusions

Polystyrene micro-and nanoplastics (PS-MNPs) are common environmental pollutants with potential respiratory health risks. Using lung cell models and organoids, we found that PS-NPs profoundly affect non-malignant lung cells, leading to increased internalization, migration, DNA damage, oxidative stress, and activation of survival pathways - hallmarks of tumor-promoting processes. In contrast, cancer cells showed only a minimal response. This study is the first to directly compare the effects of PS-NPs on malignant and non-malignant lung cells, filling a critical gap in our understanding of how microplastic exposure affects the development and progression of lung cancer. While PS-NPs did not significantly alter the behavior of cancer cells, their profound effects on non-malignant cells underscore the urgent need for further investigation into long-term lung risks associated with environmental microplastic exposure. Future research should focus on elucidating the mechanistic basis of these responses and determining the potential impact on lung disease susceptibility, progression, and treatment resistance.

## CRedit authorship contribution statement

**Büsra Ernhofer:** Conceptualization, experiments, formal analysis, investigation, validation, writing – original draft, writing – review & editing. **Anna Solta, Kristiina Boettiger, and Lisa Glatt:** Conceptualization, methodology, experiments. **Zsolt Megyesfalvi:** Conceptualization, methodology, formal analysis, supervision. **Martha Zylka:** Methodology, experiments, lung organoids investigation, validation. **Andreas Spittler, Dominik Kirchhofer, and Gerald Timelthaler:** Conceptualization, methodology, visualization, experiments. **Franziska Ferk and Miroslav Mišík:** Conceptualization, methodology, experiments, validation, resources. **Verena Kopatz:** Conceptualization, methodology, experiments. **Heinrich Kovar:** Conceptualization, methodology, experiments, lung organoids investigation, resources. **Siegfried Knasmueller:** Conceptualization, methodology, experiments, resources. **Lukas Kenner:** Methodology, resources, funding acquisition, editing. **Balazs Döme and Karin Schelch:** Conceptualization, methodology, resources, supervision, writing – review & editing, project administration, funding acquisition. We thank Barbara Dekan for technical assistance with cell culture.

All authors have reviewed, commented on the manuscript, and agreed to the submitted version.

## Data availability

Data will be made available on request.

## Acknowledgements/fundings

KS received funding from the Austrian Science Fund (FWF No. T 1062-B33). BD was supported by the Austrian Science Fund (FWF I4677). BD acknowledges funding from the “BIOSMALL” EU HORIZON-MSCA-2022-SE-01 project. BD and ZM were supported by funding from the Hungarian National Research, Development, and Innovation Office (2020-1.1.6-JÖVŐ, TKP2021-EGA-33, FK-143751 and FK-147045). ZM was supported by the New National Excellence Program of the Ministry for Innovation and Technology of Hungary (UNKP-20-3, UNKP-21-3 and UNKP-23-5), and by the Bolyai Research Scholarship of the Hungarian Academy of Sciences. ZM is also the recipient of the International Association for the Study of Lung Cancer/International Lung Cancer Foundation Young Investigator Grant (2022). VK and LK are supported from MicroONE, a COMET Modul under the lead of CBmed GmbH, which is funded by the federal ministries BMK and BMDW, the provinces of Styria and Vienna, and managed by the Austrian Research Promotion Agency (FFG) within the COMET—Competence Centers for Excellent Technologies—program. Financial support was also received from the Austrian Federal Ministry of Science, Research and Economy, the National Foundation for Research, Technology and Development, and the Christian Doppler Research Association, as well as Siemens Healthineers for their financial and scientific support. L.K. was also supported by a European Union Horizon 2020 Marie Sklodowska-Curie Doctoral Network grants (FANTOM, n. P101072735 and eRaDicate, n. 101119427) the Christian-Doppler Lab for Applied Metabolomics (CDL-AM), and the Austrian Science Fund (grants FWF: P26011, P29251, P34781 as well as the International PhD Program in Translational Oncology IPPTO 59.doc.funds). Additionally, this research was funded by the Vienna Science and Technology Fund (WWTF), grant number LS19-018. L.K. is a member of the European Research Initiative for ALK-Related Malignancies (www.erialcl.net). HK was supported by the Austrian Science Fund (FWF P35353-B).

