## Supplementary for "Small Particles, Big Problems: Polystyrene nanoparticles induce DNA damage, oxidative stress, migration, and mitogenic pathways predominantly in non-malignant lung cells"

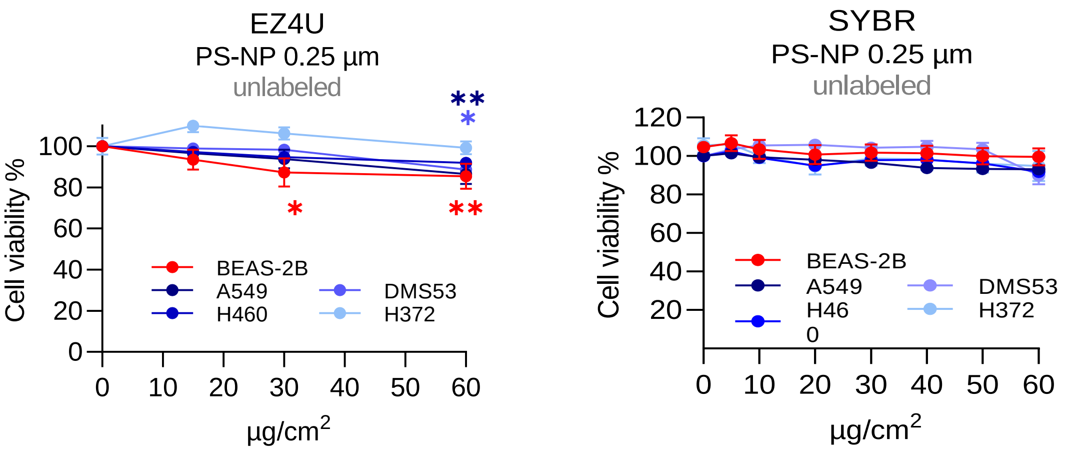


**Supplementary Figure S1: Impact of PS-NPs on variety of lung-derived cell lines.** Dose response curves of all lung cell lines after exposure to unlabeled 0.25 µm PS-NPs for 72 h with doses ranging from 15 to 60 µg/cm². Cell viability was performed using EZ4U and SYBR assays. Data are shown as mean ± SEM of three independent experiments performed in triplicates. ANOVA followed by Dunnett's multiple comparisons test. *p ≤ 0.05 and **p ≤ 0.01.

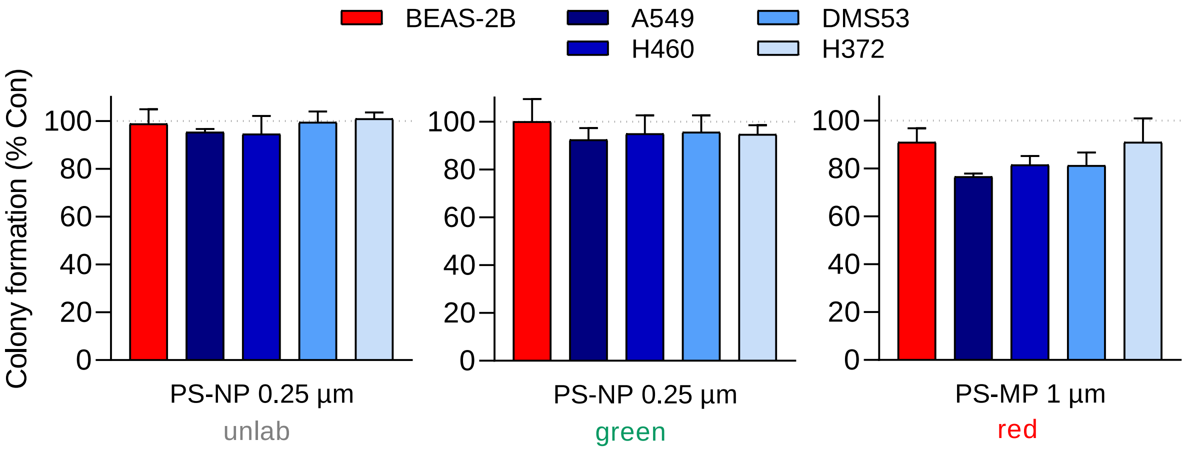


**Supplementary Figure S2: Impact of colony formation in a variety of lung-derived cell lines.** PS-MNP treatment showed no effect on cell viability in long-term. Quantification of colony formation assay for all cell lines after a 10-day incubation with 10 µg/cm² PS-MNPs. Data are shown as mean ± SEM of three independent experiments performed in triplicates.

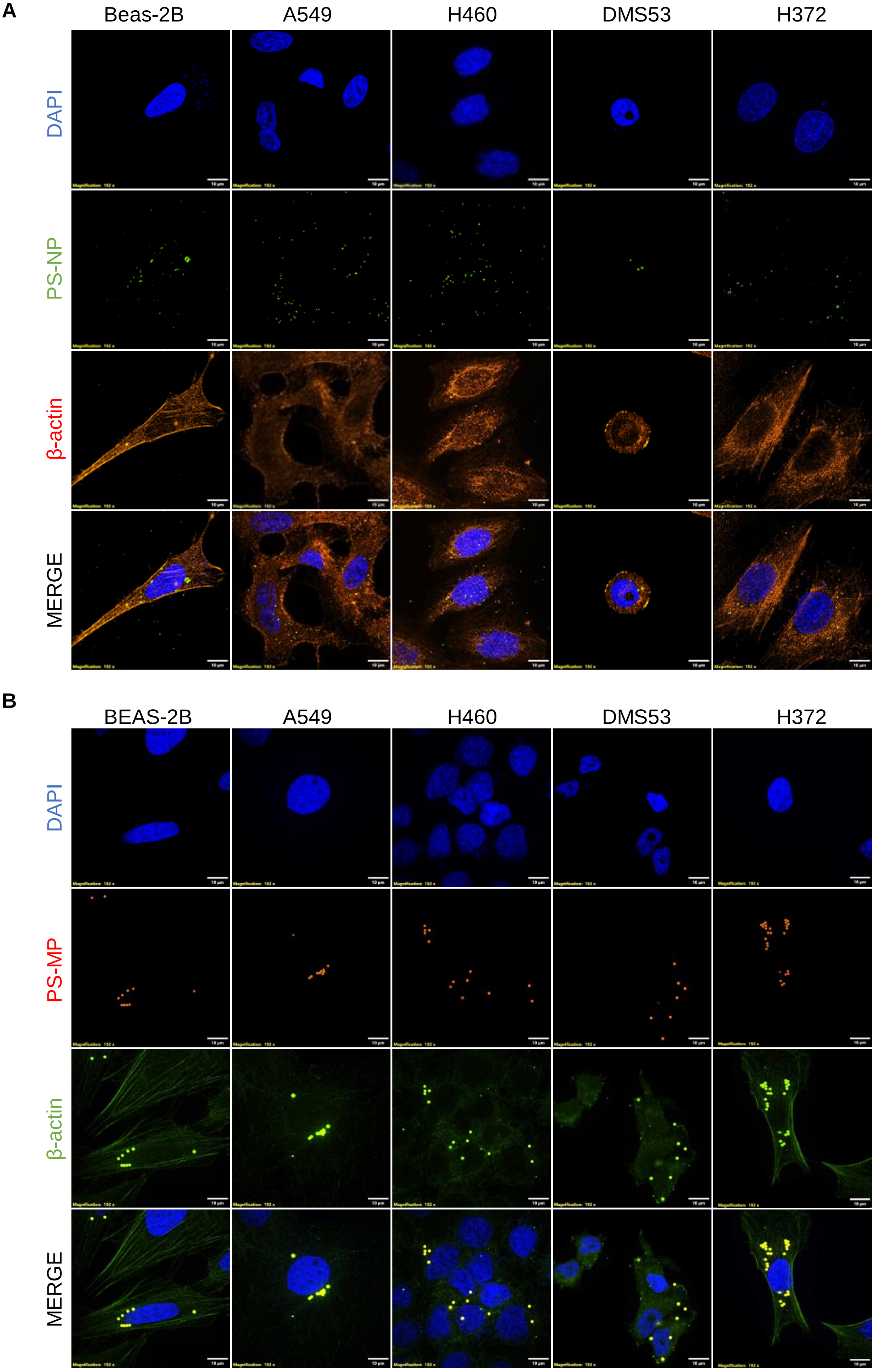


Ernhofer et al. – Supplement Figure S3

**Supplementary Figure S3: PS-MNP uptake in all lung cell lines. A)** and **B)** Representative confocal microscopy images showing internalization of PS-MNPs (24 h, 1 µg/cm$^2$) by non-malignant (BEAS-2B) and malignant (A549, H460, DMS53, and H372) lung cells. PS-NPs (0.25 µm) are shown in green and PS-MPs (1 µm) in red. Corresponding β-actin was labeled either in red or green. Yellow indicates the merged double-positive fluorescence signal of 1 µm PS-MPs, which is also observed in the green channel. Nuclei were stained with DAPI (blue). Scale bar: 10 µm.

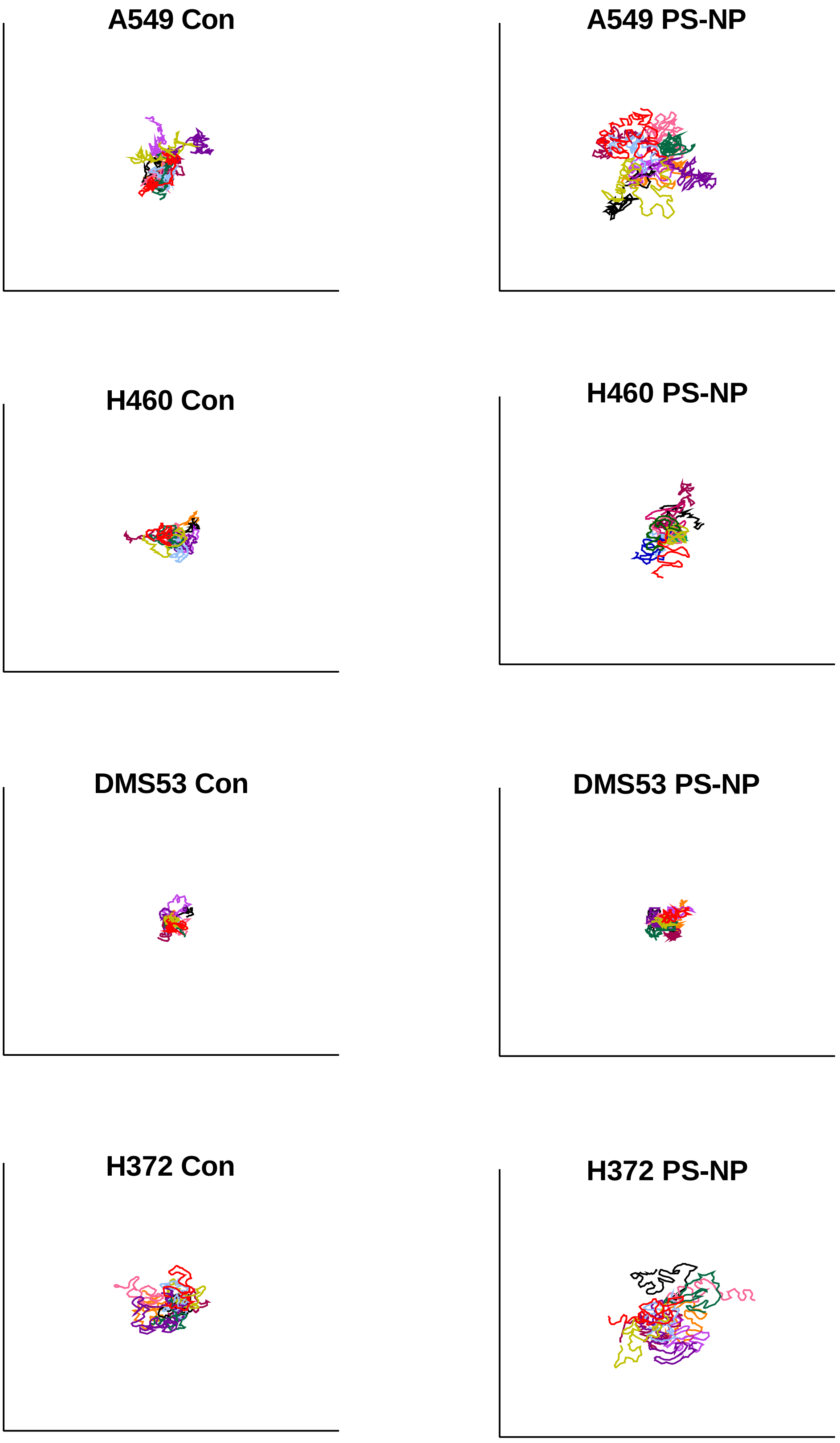


**Supplementary Figure S4: Migrated distance of lung cells after 96 h exposure to 10 µg/cm² PS-NPs.** Origin plots display 10 representative cell tracks relative to one origin (0/0). Lines in color represent individual cells.

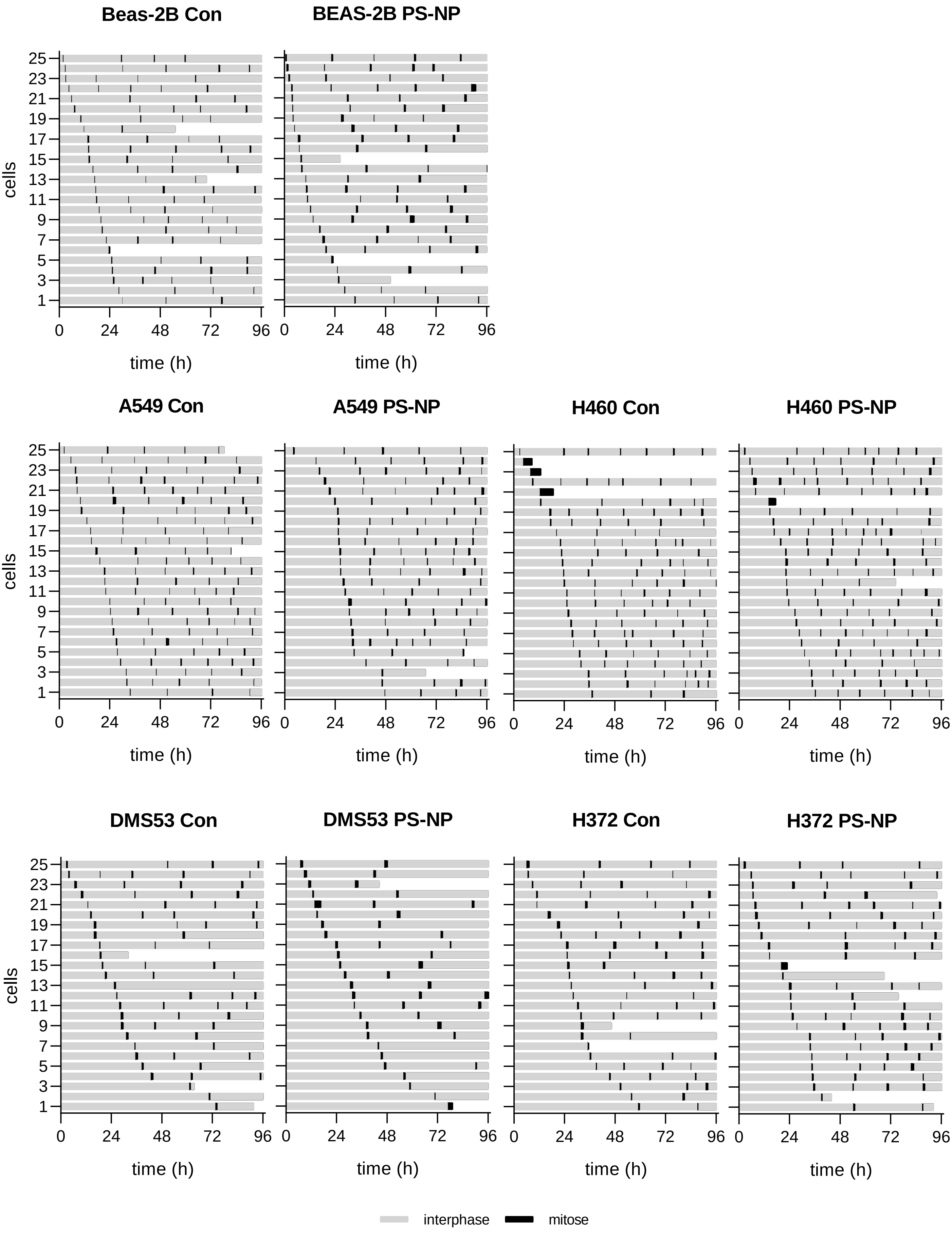


Ernhofer et al. – Supplement Figure S5

**Supplementary Figure S5: Cell fate map following PS-NP treatment revealed no changes in cell division and growth behavior.** Cells were monitored via videomicroscopy over 96 h of exposure to 10 µg/cm² unlabeled PS-NP, with images captured at 10 min intervals. The gray and black colored bars indicate the interphase and M-phase, respectively. Bars that do not reach the end of the graph represent cells that died earlier. Each bar represents an individual cell.

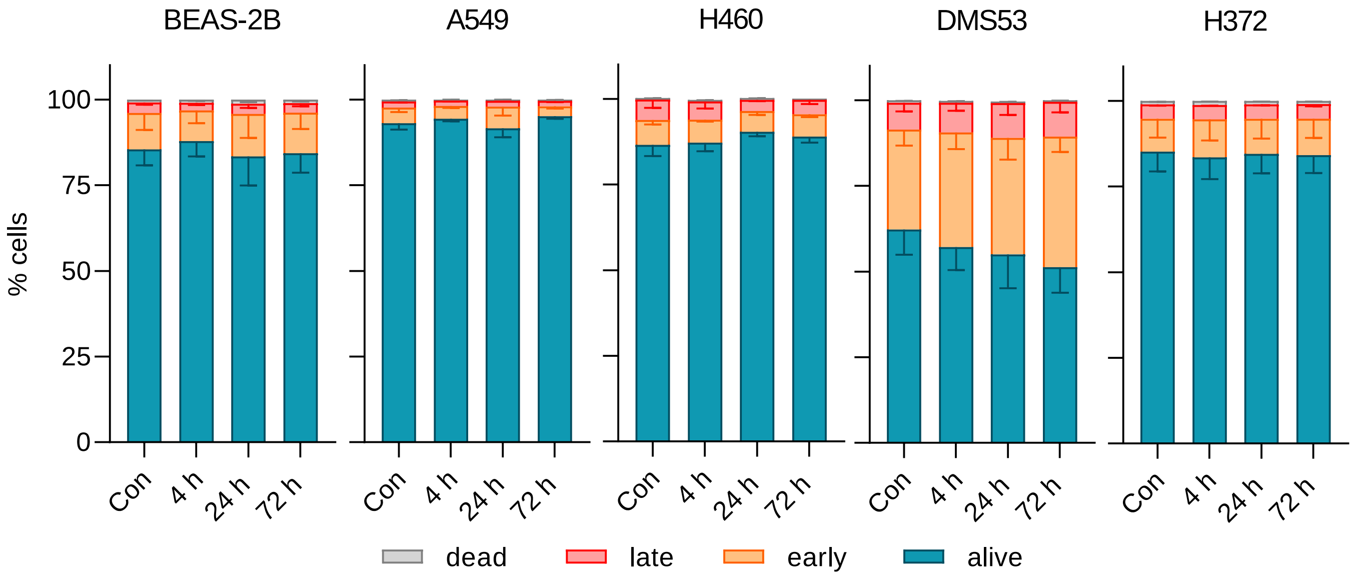


**Supplementary Figure S6: Lung cell lines showed no differential response to PS-MNP treatment.** Flow-cytometry-based apoptosis assay using PI and Annexin V staining was conducted at different time points with 10 µg/cm² of unlabeled PS-NPs. Data are shown as mean ± SEM of three independent repeats performed in triplicates (blue – live, orange – early, red – late, gray – necrotic cell fractions).

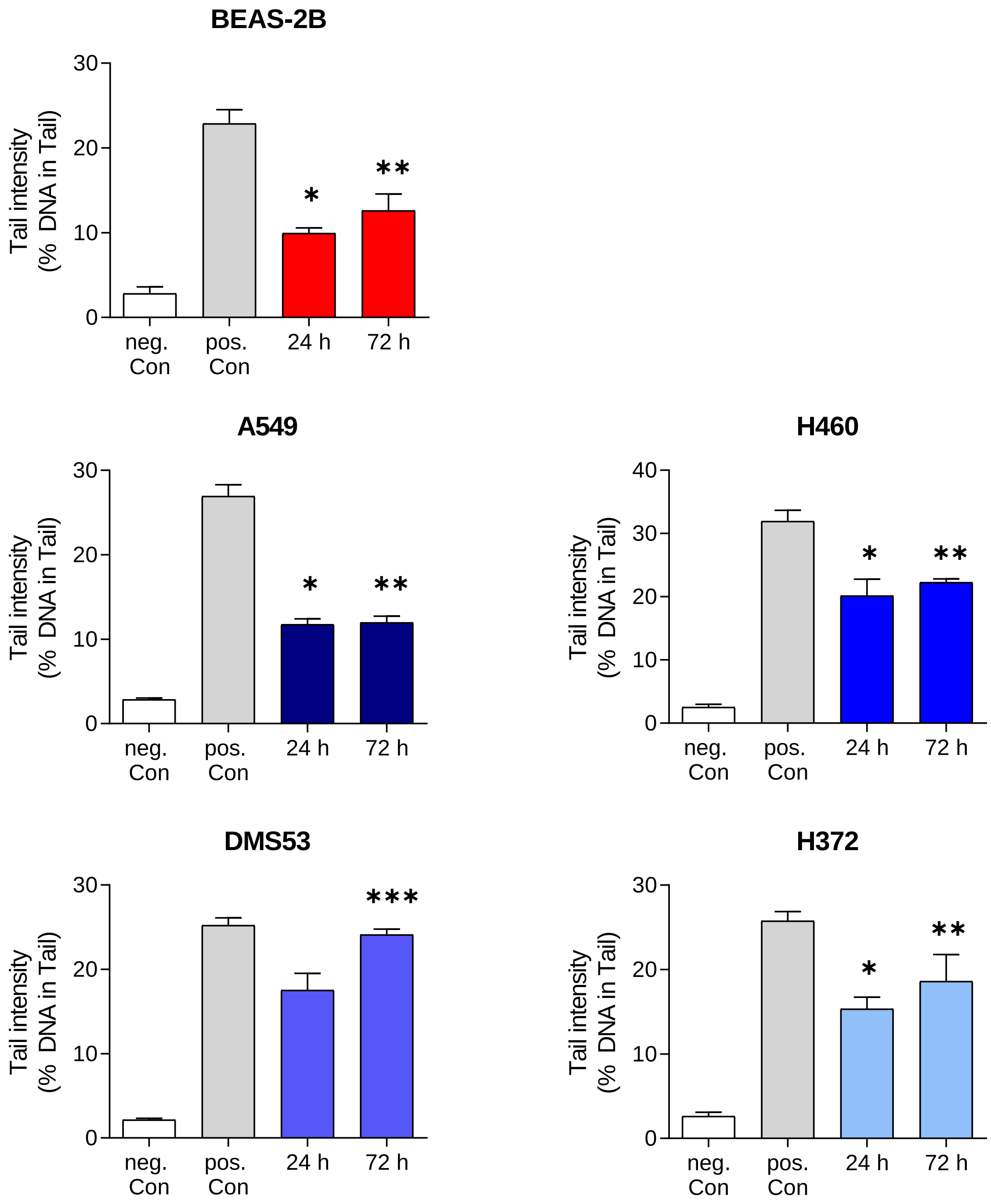


**Supplementary Figure S7: Single DNA damage graphs of non-malignant and malignant lung cell lines.** Standard comet assays revealed DNA damage in all lung cells. Baseline DNA damage levels after 24 h and 72 h of exposure to 10 µg/cm² unlabeled PS-NP are expressed as a percentage of DNA intensity in tail compared to untreated control cells. Negative (PBS) and positive ($H_2O_2$) controls were included for each cell line. Data are shown as mean ± SEM of two independent experiments. ANOVA followed by Dunn's multiple comparisons test. *$p \leq 0.05$, **$p \leq 0.01$, ***$p \leq$. 0.001.

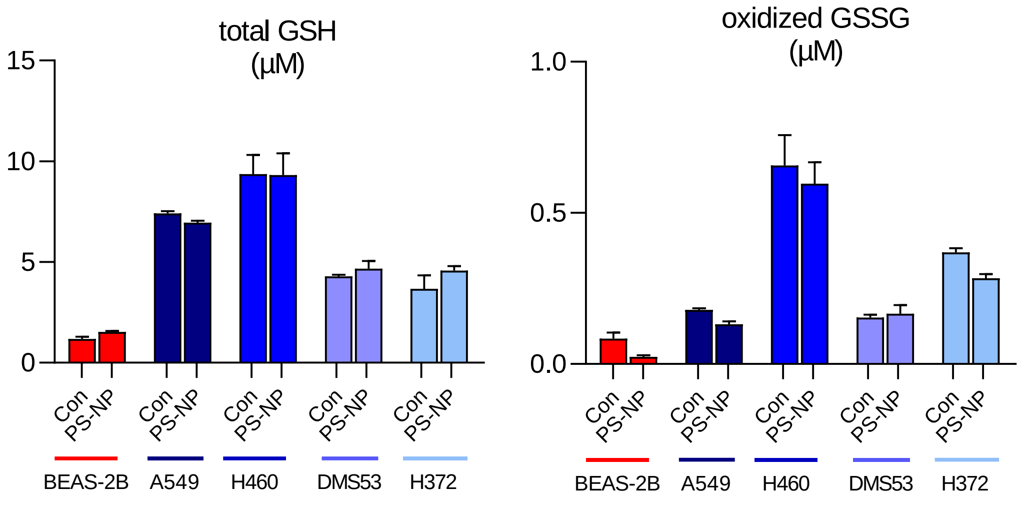


**Supplementary Figure S8: Quantification of total and oxidized glutathione.** Total glutathione (GSH + GSSG) and oxidized glutathione (GSSG) levels (µM) of all cell lines were determined using a luminescence-based GSH/GSSG-Glo assay (Promega). Cells were treated with unlabeled PS-NP (10 µg/cm², 24 h) and data are the mean ± SEM of at least two independent experiments. ANOVA and Tukey's multiple comparisons test. *p ≤ 0.05, **p ≤ 0.01, and ***p ≤ 0.001 compared to non-treated control cells, respectively.

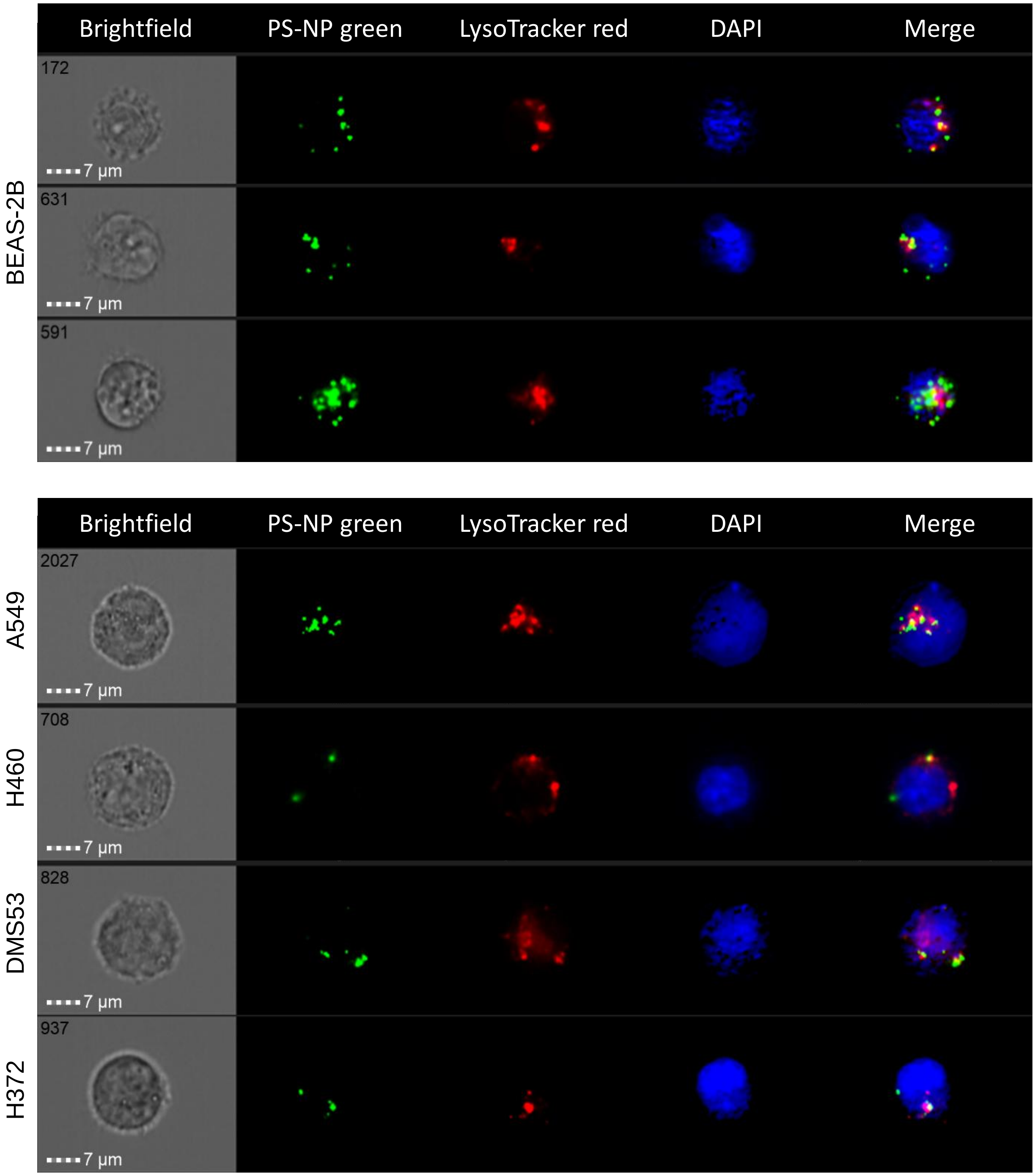


**Supplementary Figure S9: Co-localization with lysosomes and 0.25 µm PS-NP.** Representative images showed lysosomes (LysoTracker, red) and PS-NPs (green, 10 µg/cm²) after 24 h incubation in all lung cells, acquired by AMNIS flow cytometry. Upper row: non-malignant (BEAS-2B) and bottom row: malignant (A549, H460, DMS53, and H372). Scale bar: 7 µm.

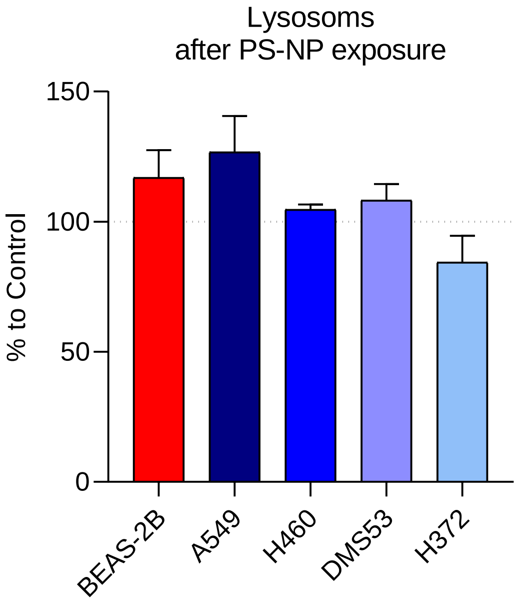


**Supplementary Figure S10: Quantification of lysosomes.** Flow-cytometry-based quantification of lysosomes using LysoTracker (Red) after 24 h treatment to 0.25 µm unlabeled PS-NP (10 µg/cm$^2$) in all lung cell lines. Data shown as mean ± SEM of three independent experiments and compared to non-treated control cells, respectively.

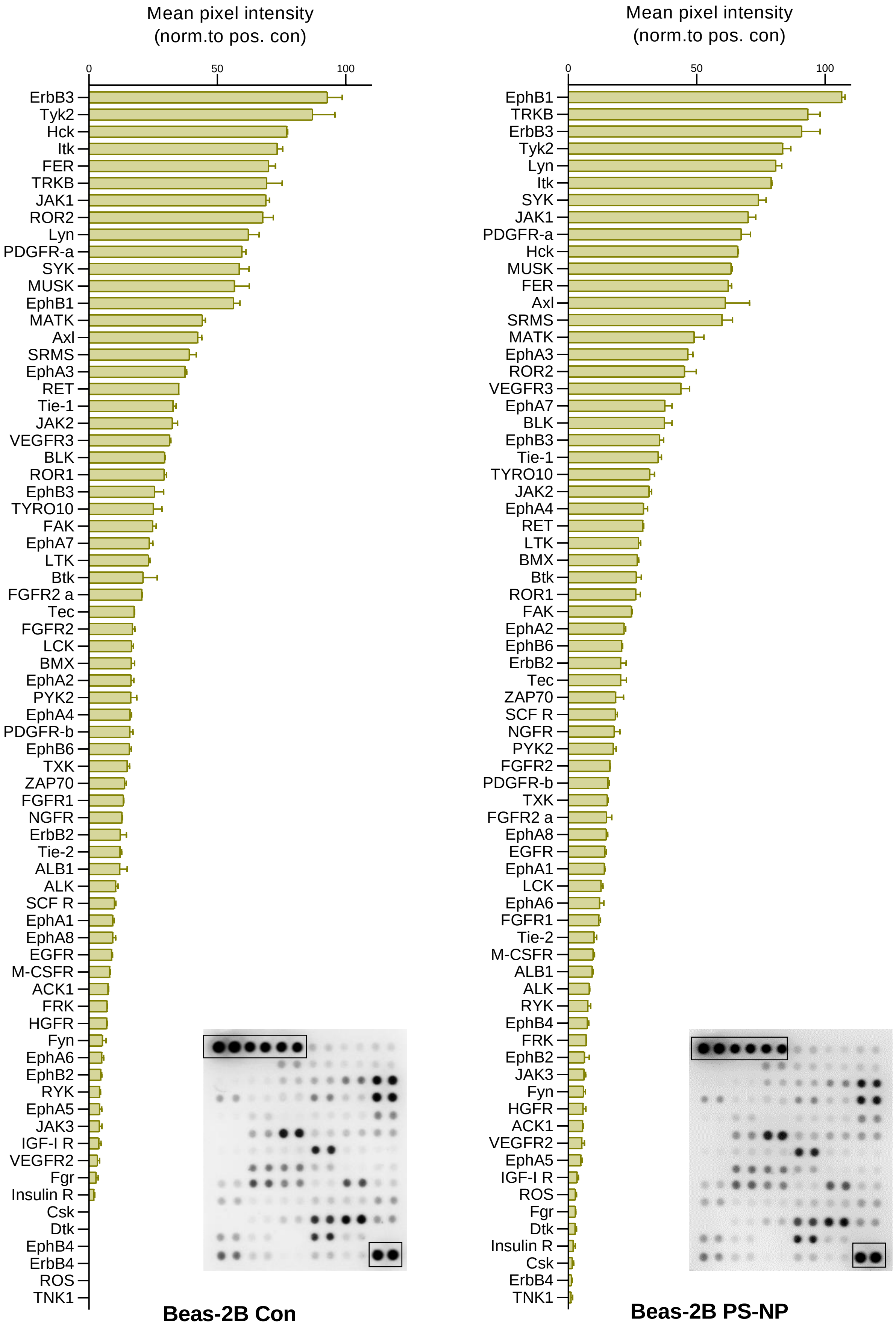


**Supplementary Figure S11: RTK results of untreated and treated BEAS-2B with PS-NP for 24 h.** Results and representative images of RTK membrane. Data are shown as mean ± SEM and positive controls of each blot was used as reference (rectangles).
